# An ultra-long heavy chain bovine antibody neutralizes SARS-CoV-2 and reacts broadly with sarbecoviruses

**DOI:** 10.1101/2025.01.03.631215

**Authors:** Theocharis Tsoleridis, Chengcheng Fan, Emily J. Park, Joshua D. Duncan, Parul Sharma, Sophie Wartnaby, Joseph G. Chappell, Anja Kipar, Eleanor G. Bentley, Adam Kirby, Ximeng Han, Christopher M. Coleman, Igor Santos, Dalan Bailey, Andrew I. Flyak, C Patrick McClure, Semi Rho, Daniel X. Johansson, Mats A.A. Persson, Alex W. Tarr, David Haig, James P. Stewart, Pamela J. Bjorkman, Richard A. Urbanowicz, Jonathan K. Ball

**Author notes:** Corresponding authors. (TT); (RAU); (JKB). These authors contributed equally to this work (Theocharis Tsoleridis, Chengcheng Fan, Emily J. Park). Present address: Centre for Human Genetics, Nuffield Department of Medicine, The University of Oxford; Oxford, UK. Present address: Jenner Institute, University of Oxford; Oxford, OX3 7DQ, UK.

## Abstract

The ongoing threat of new SARS-CoV-2 variants and other sarbecoviruses has driven efforts to develop broadly neutralizing monoclonal antibodies (mAbs). This study used immunized cattle, known for producing antibodies with ultra-long CDRH3 domains, to generate 33 mAbs, ten of which had ultra-long CDRH3s (>50 amino acids). Of these, mAbs P7 and 99, demonstrated broad and potent neutralization. Notably, mAb P7 neutralized all tested variants, including SARS-CoV-1, with IC_50_ values between 0.05 and 9.2 µg/mL, and showed cross-reactivity with RBDs from various sarbecoviruses. Structural analysis revealed that mAb 99 binds the spike protein’s RBD at the ACE2 binding site. Although the exact binding of P7 wasn’t resolved, evidence suggests it targets a hidden epitope, promoting spike trimer dissociation via its extended CDRH3. In Syrian hamsters, both mAbs significantly reduced lung viral loads. These results support the potential of bovine-derived mAbs, particularly those with ultra-long CDRH3s, for future antiviral therapies.

## INTRODUCTION

Since its emergence in 2019, SARS-CoV-2, the causative agent of COVID-19, has spread worldwide and has claimed more than 6.9 million lives [https://data.who.int/dashboards/covid19/cases]. Adaptive immunity (both neutralising antibodies (nAbs) and T cell-mediated immunity) plays a crucial role in protection against severe disease and clearing virus (*1, 2*), and this is induced following natural infection and immunisation (*3–5*). The introduction of vaccines against SARS-CoV-2 was a success, preventing 14 million associated deaths in the first year of the implementation of vaccination programmes globally (*6*). The majority of licenced vaccines for use in humans were developed using the ancestral Lineage A (Wuhan) SARS-CoV-2 spike (S) protein (*7*). However, the constant evolution of the virus has given rise to new variants of concern (VOCs), such as the Omicron variants, which have the potential to escape from antibodies in individuals with prior exposure to the virus and/or vaccination (*8*). Despite vaccination, immunocompromised patients often show impaired responses and develop severe disease, highlighting the need for therapeutic monoclonal antibodies with potent cross-reactivity (*9*). In addition, human cell tropism reported for many species within the *Sarbecovirus* subgenus highlights their potential for future spill-over events (*10*). Therefore, the development of broad-acting therapeutic interventions remains a high priority.

Monoclonal antibodies (mAbs) against SARS-CoV-2 have been crucial in the development of effective treatment options against COVID-19. Broadly neutralising antibodies have been isolated from individuals who had recovered from COVID-19 (*11, 12*) and from mice (*13, 14*). The first therapeutic mAb clinical trials started a few months after the release of the virus genome (*15*). The use of therapeutic mAbs remains important for the treatment of patients with severe disease and vulnerable populations such as elderly and immunocompromised individuals. However, the emergence of new variants, such as Omicron and its sublineages, has reduced the efficacy of the currently licenced therapeutic mAbs (*16–18*). Hence, the continued development and discovery of effective therapeutic mAbs against current and potential future variants remains essential.

Bovines are known to produce antibodies with unique features, including very long heavy chain complementary determining region 3s (CDRH3s) averaging 26 amino acids and a subset of’ultra-long’ (UL) antibodies with CDRH3 lengths of up to 75 amino acids (*19, 20*). An UL-CDRH3 forms a distinct extended conformation consisting of a stalk region and a cysteine-rich knob domain that mediates antibody-antigen complex formation (reviewed in (*21*)). Given the extended conformation of the UL-CDRH3 domain, it has been proposed that these antibodies could target epitopes of viral glycoproteins that are otherwise inaccessible to human B cell repertoires (*22*). Previous studies have demonstrated that immunisation of *Bos taurus* with soluble HIV Env trimers can elicit UL-CDRH3 broadly neutralising antibodies when tested against large panels of HIV isolates (*23–25*). Furthermore, surface display libraries generated from naïve *B. taurus* B-cells have shown that anti-SARS-CoV-2 spike paratopes exist within this B cell receptor (BCR) repertoire (*26*). In this current study, we serially immunised cattle with different forms of the SARS-CoV-2 spike protein and, through a combination of single B cell sorting and phage display approaches, isolated and characterised a panel of bovine mAbs. Two mAbs exhibited potent and broad neutralising activity across a range of major variants of concern, and one of these, a mAb with an ultra-long 61 residue CDRH3, exhibited high affinity binding to the receptor-binding domains (RBDs) of diverse sarbecoviruses. This study demonstrated that *B. taurus* can be utilised as a source of novel broad-acting therapeutic mAbs to mitigate the potential impacts of future coronavirus spillover events.

## RESULTS

### SARS-CoV-2 immunised cattle produce potent and broadly neutralising antibodies

To investigate bovine immune responses against SARS-CoV-2 spike, a series of immunisations were performed on two Holstein bullocks. The immunisation process lasted for 242 days and a total of five immunisations were performed: one prime immunisation and two boosts with the S1 subunit of Lineage A S1 spike (aa 1-674), one boost with Lineage A spike ectodomain (S1 and partial S2) and a final boost with Lineage A trimeric spike, (transmembrane domain replaced with a trimerization domain) (Fig. 1**A**). Sera and blood were collected 4-and 8-days after each boost. Sera binding was measured using an antigen binding assay utilising spikes from different variants as target antigen. Maximum breadth and potency were observed in sera harvested at D242 post-immunisation (data not shown). Neutralisation assays using lentivirus-based pseudotypes supplemented with spikes representative of the original Lineage A SARS-CoV-2, variants of concern (Alpha, Beta, Gamma, Delta, Omicron variants B.1.1.529 and BA.5), and to test pan-species breadth, SARS-CoV-1 Frankfurt 1 strain yielded ID_50_ dilution values for Lineage A, Alpha, Beta, Omicron and Omicron BA5 of approximately 1:1000, and for Gamma, Delta and SARS-CoV-Fra1 1:100 (Fig. 1**B**).

**Figure 1.**
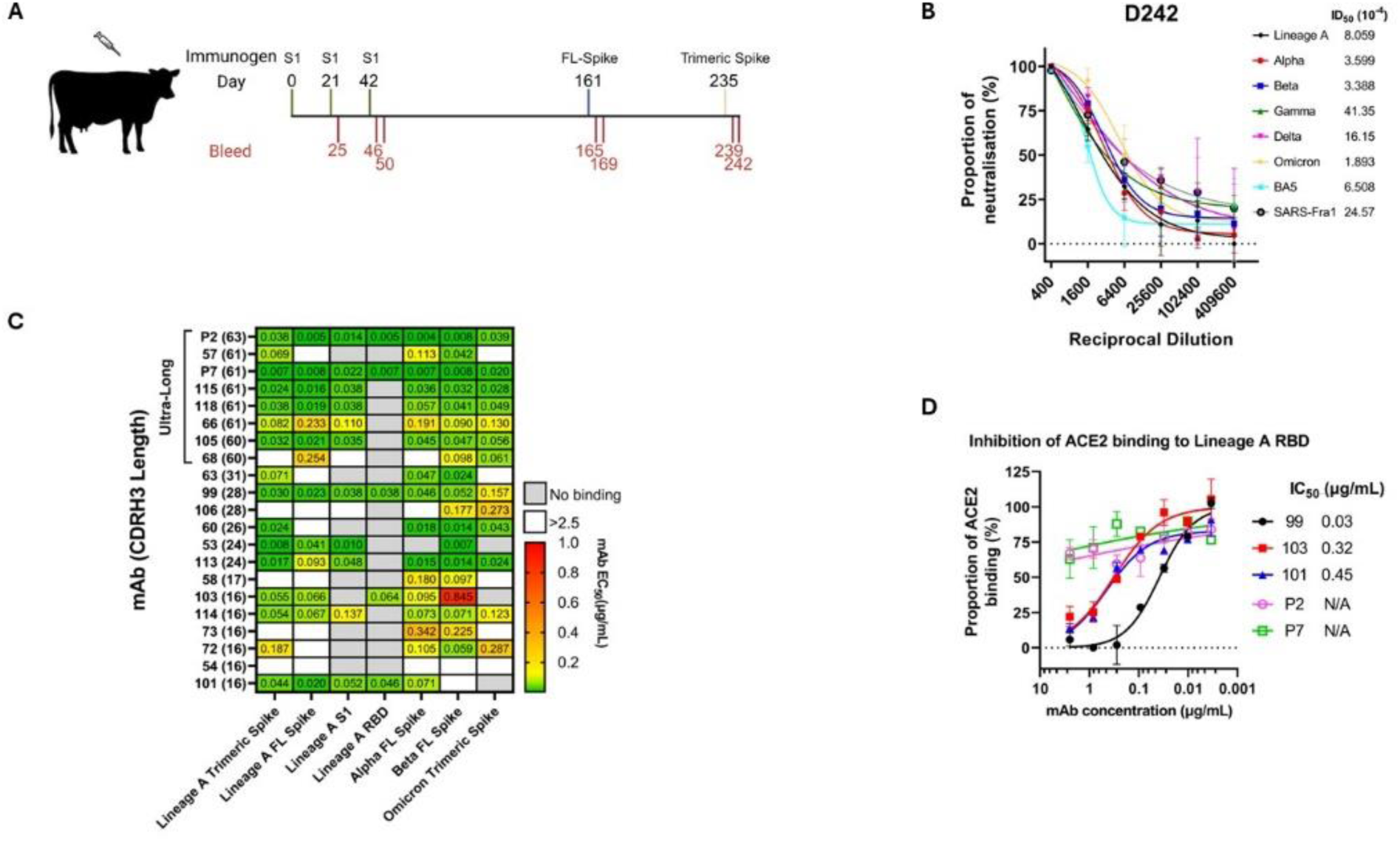
(**A**) Schematic representation of the immunisation schedule. The process lasted for 242 days and it consisted of one prime immunisation with Lineage A S1 spike, two boosts with the same antigen, one boost with Lineage A full length spike and one boost with Lineage A trimeric spike. Blood was collected 4 and ∼8 days post immunisation. (**B**) Neutralisation assay using cow serum from the final timepoint (D242) against pseudotyped SARS-CoV-2 variants and the original SARS-CoV. The serum was able to neutralise all pseudotypes tested and ID_50_ values were calculated. (**C**) Table showing the EC_50_ values (µg/mL) of spike binding mAbs against different forms of Lineage A spike (trimeric, full length, S1, RBD) and other variant spikes (Alpha, Beta, Omicron (BA.5)). The mAbs are divided in ultra-longs and non-ultra-longs based on their CDRH3 length in amino-acids. (**D**) Competition ELISA of the five RBD-binding mAbs (P2, P7, 99, 103, 101) with human ACE2 against Lineage A RBD. Antibodies 99, 103 and 101 showed inhibition of ACE2 with IC_50_ values of 0.03, 0.32 and 0.45 µg/mL, respectively. Antibodies P2 and P7 showed partial inhibition. Mean and SD shown on graphs

Having established the presence of potent and broadly neutralising serum antibody responses at D242 from both animals, we then went on to generate a panel of spike-specific monoclonal antibodies. To achieve this, two methods were performed. The first utilised single-cell sorting of antigen-specific B cells in which peripheral blood mononuclear cells (PBMCs) from D242 were subjected to flow cytometry using a dead/live gate and Lineage A trimeric spike labelled with two different fluorochromes (fig. S1). Single B-cells specific for Lineage A trimer were sorted into 96-well plates and then PCR amplified using an IgG-specific nested primer PCR to retrieve individual heavy and light chain pairings. These were cloned into a human IgG1 expression cassette and expressed in mammalian cells. This yielded an initial panel of 31 candidate mAbs. The second method used a phage display approach in which antigen-specific B cells from D242 were isolated by flow cytometry using dual-staining with soluble 2P and GSAS substituted furin cleavage site trimeric spike protein. Isolated cells were pooled, RNA extracted, reverse transcribed and the resulting cDNA used as template in a nested primer PCR designed to preferentially amplify heavy chains derived from the VH1-7 germline, from which all ultra-long antibodies derive (*27*). Resulting PCR products were cloned, paired with a common light chain to generate Fab-p3-phage display libraries, which were then biopanned against SARS-CoV-2 Lineage A trimer followed by a second round of biopanning against Omicron BA.5 trimeric spike. This method identified a further two mAbs, P2 from round one and P7 from round 2, which were subcloned into a human IgG1 expression cassette and expressed in mammalian cells and added to the panel (table S1). The length of the CDRH3 of each mAb in the panel ranged between 16-63 amino acids (aa). Ten of 33 antibodies were ultra-long with CDRH3 >50aa and they all encoded the same V-gene segment, VH1-7.

### Isolated monoclonals show pan-sarbecovirus reactivity *in vitro*

In the next step, we characterised the reactivity and neutralising ability of the mAb panel. Twenty-one antibodies showed reactivity to Lineage A spike protein, eight of which were ultra-long (fig. S2) and 13 were non-ultra-long (fig S3). Nine mAbs bound to all variant spike proteins. Moreover, five antibodies were identified as RBD binders (99, 101, 103, P2 and P7) whereas the other mAbs bound outside the RBD (Fig. 1**C**). A competition ELISA assay was performed on the five RBD binding antibodies to assess whether they inhibit ACE2 receptor binding to Lineage A RBD spike. Antibody 99 showed the strongest inhibition of ACE2 (IC_50_ of 0.03 µg/mL), followed by antibodies 103 and 101 (IC_50_ 0.32 µg/mL and 0.45 µg/mL, respectively). Antibodies P2 and P7 showed partial inhibition with a plateau of 70% ACE2 binding, suggesting accessibility hinderance to the RBD (Fig. 1**D**).

Having characterised the binding properties of each mAb in the panel, the next step was to assess their neutralisation potencies and breadth. To achieve this, the entire mAb panel was used in single-point neutralisation assays (1 µg/mL final mAb concentration) performed against pseudotyped SARS-CoV-2 viruses, including Lineage A, Alpha, Beta, Gamma, Delta and Omicron (B.1.1.529). Antibodies that were able to neutralise >75% infectivity of all variants were selected for further in-depth investigation. These criteria were fulfilled by four UL antibodies (P2, P7, 105, 118) and one non-UL (99) (Fig. 2**A**), the latter showing 100% neutralisation against all variants at 1 µg/mL.

**Figure 2.**
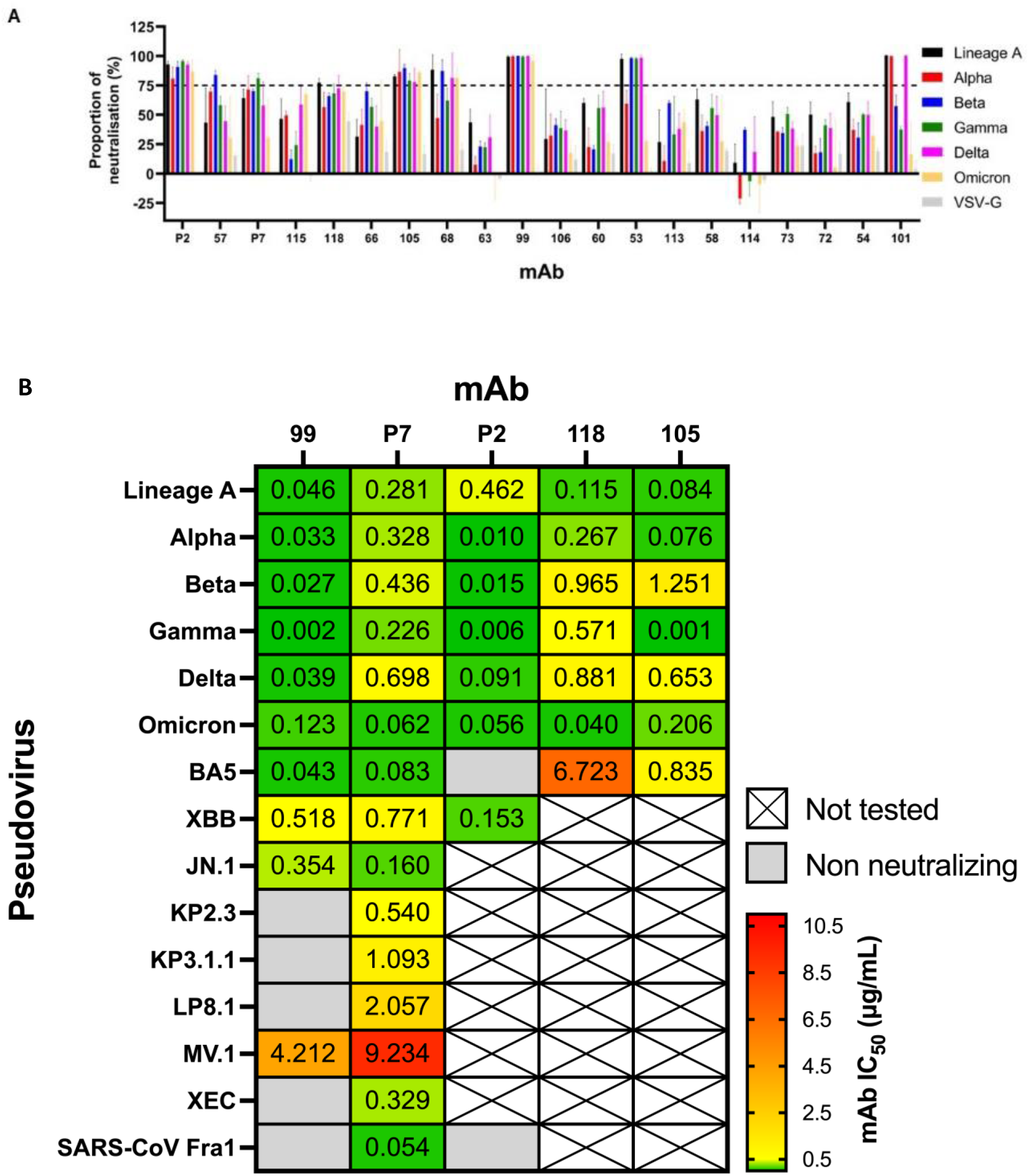
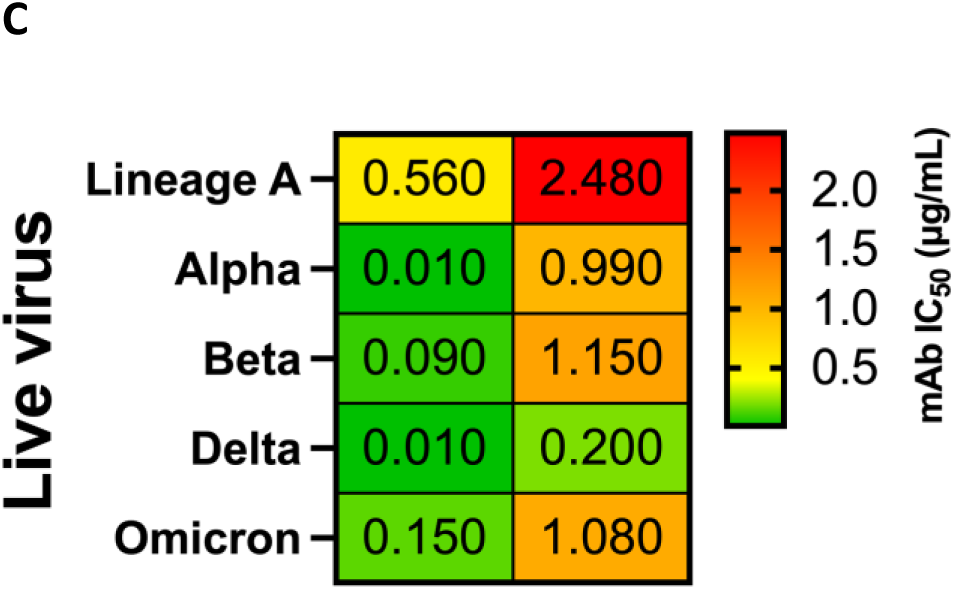
(**A**) Single point neutralisations of the mAb panel against SARS-CoV-2 pseudotypes (Lineage A, Alpha, Beta, Gamma, Delta and Omicron). The antibodies were divided into ultra-longs and non-ultra-longs according to their CDRH3 length in amino-acid. Antibodies able to neutralise all variants at 75% minimum were considered broadly neutralising and were investigated further. These antibodies were P2, P7, 105, 118 and 99. (**B**) Table with IC_50_ values (µg/mL) for the candidate mAbs against different SARS-CoV-2 pseudotypes (Lineage A, Alpha, Beta, Gamma, Delta, Omicron, BA5, XBB, JN.1, KP2.3, KP3.1.1, LP8.1, MV.1, XEC) and SARS-CoV-1. Antibody P7 neutralised all pseudotypes, including SARS-CoV-1. The second most broad neutralising antibody was 99, which neutralised 10 SARS-CoV-2 pseudotypes but not SARS-CoV-1. NN stands for “not neutralising” and N/A means that no experiments were performed for this mAb/pseudotype combination. (**C**) Table showing IC_50_ values (µg/mL) for antibodies P7 and 99 against SARS-CoV-2 live virus (Lineage A, Alpha, Beta, Delta, Omicron). Both antibodies neutralised every variant; however, mAb 99 exhibited consistently lower IC_50_ values against all variants.

These five candidate mAbs were subjected to further neutralisation experiments against pseudotyped SARS-CoV-2 viruses to determine IC_50_ values against different variants of concern (Fig. 2**B**). The non-RBD UL antibodies 105 and 118 had a similar breadth across the variants up to and including Omicron, but 118 had a much higher IC50 against Omicron BA.5 (fig. S4). The IC_50_ values for mAb 105 were between 0.001 µg/mL (Gamma) and 0.84 µg/mL (Omicorn BA.5) and for mAb 118 between 0.04 µg/mL (Omicron) and 6.7 µg/mL (Omicron BA.5). mAb P2 had good potency to historic SARS-CoV-2 variants with IC_50_ values ranging from 0.006 µg/mL (Gamma) to 0.46 µg/mL (Lineage A); however, did not neutralise variant Omicron BA.5 and SARS-CoV-1 (fig. S4). mAb 99 performed consistently well against historic SARS-CoV-2 variants, including variants such as Omicron BA.5 and XBB, with IC_50_ values ranging from 0.002 µg/mL (Gamma) to 0.52 µg/mL (XBB) (fig. **S4**). It was able to neutralise JN.1 (ID_50_ of 0.35 µg/mL) and showed limited potency against MV.1 (ID_50_ of 4.21 µg/mL) but was unable to neutralise more contemporary strains (KP2.3, KP3.1.1, LP8.1 and XEC). The antibody with the greatest neutralising breadth was P7, which was able to neutralise all variants tested, including the 2002 epidemic SARS-CoV-1 strain, although potency was limited against MV.1 (ID_50_ of 9.23 µg/mL) with IC_50_ values ranging from 0.05 µg/mL (SARS-CoV-1) to 9.23 µg/mL (MV.1) (fig. S4).

The two neutralising antibodies showing maximal potency and breadth, P7 and 99, were also tested against Lineage A, Alpha, Beta, Delta and Omicron variants in an authentic virus neutralisation assay. Both mAbs showed neutralising activity against all variants. The IC_50_ values for P7 ranged between 0.2 µg/mL (Delta) and 2.48 µg/mL (Lineage A), whereas for mAb 99, values ranged between 0.01 µg/mL (Alpha and Delta) and 0.56 µg/mL (Lineage A) (Fig. 2**C**, fig. **S5**). Interestingly, both antibodies performed better against variants of concern (especially Delta) than against the original Lineage A, which the cattle were immunised with. Comparison between the pseudotype and the authentic virus assay showed similar IC_50_ values for 99, except for Lineage A which was less potent in the authentic virus assay. However, there was disparity between the two assays for P7, which consistently showed reduced neutralisation activity against all authentic virus variants tested.

To determine the full breadth of the RBD-binding mAbs (P2, P7, 99, 103 and 101), binding assays were performed on a panel of 23 SARS-CoV-2 and other sarbecovirus RBD spike regions. Remarkably, P7 showed strong RBD binding across the panel, with EC_50_ values ranging from 0.083 µg/mL to 0.311 µg/mL, suggesting targeting of a conserved epitope within the RBD (Fig. **3**). The second most broadly reactive antibody was mAb 99, followed by P2, binding 8 of 23 and 7 of 23 RBDs, respectively. Antibodies 103 and 101 were the least broad, binding 4 of 23 and 3 of 23 RBDs, respectively, and they showed similar binding patterns; all of the RBDs they bound were also bound by mAb 99 (fig. S**6**).

**Figure 3.**
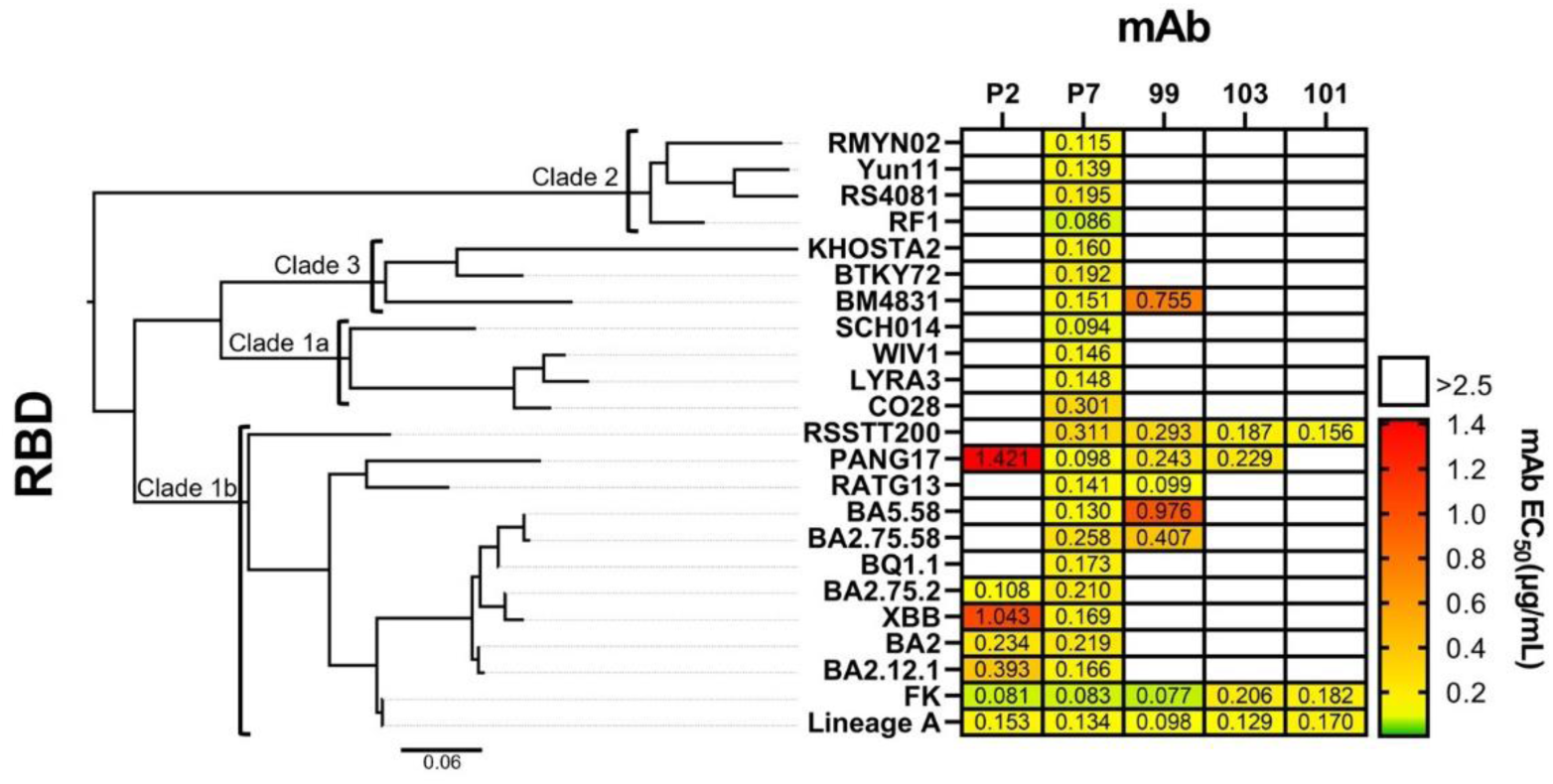
EC_50_ values (µg/mL) of the five RBD-binding mAbs against a panel of 23 SARS-CoV-2 variants and other sarbecovirus RBDs. Any EC_50_ >2.5 µg/mL was disregarded. The most broadly reactive antibody was P7, which bound every RBD in the panel, with EC_50_ values ranging from 0.094 to 0.311 µg/mL. The rest of the antibodies bound 8 of 23, 7 of 23, 4 of 23 and 3 of 23 for 99, P2, 103 and 101, respectively. Next to the table is a phylogenetic tree of the corresponding sarbecovirus RBDs

### mAbs 99 and P7 protect in *in vivo* challenge model

To assess the efficacy of candidate mAbs *in vivo*, we performed authentic virus challenge experiments in Syrian hamsters. The study consisted of four groups of six animals that were challenged with the SARS-CoV-2 Delta (1 × 10^4^ pfu) on day 0, and then treated with mAb or placebo on day 1, with weight changes followed over 7 days. The animals in the untreated group showed a significant and progressive weight loss (10% by day 7), whereas all animals treated therapeutically showed only a small weight loss 24 h after viral challenge, and from day 2 had recovered to at least pre-challenged weights (Fig. 4**A**). The weight loss in all mAb-treated groups was significantly different from the control group given PBS alone (p < 0.05 and p < 0.01; repeated measures two-way ANOVA with Dunnett’s multiple comparison test). Analysis of viral load in the lungs at day 7 by qPCR for Nucleoprotein (N1) RNA showed a decrease in the mean value in treated compared to the untreated control animals (Fig. 4**B**). This decrease was significantly different between mAb 99 (2×10^3^ copies of N1/μg of RNA; p = 0.0006), P7 (3×10^3^ copies of N1/μg of RNA; p = 0.0008) and PBS only (4×10^4^ copies of N1/μg of RNA). Similar results were observed for the commercial control Sotrovimab (7×10^2^ copies of N1/μg of RNA; p = 0.0006; one-way ANOVA with Dunnett’s multiple comparison). The measured viral load in the nasal tissue was also significantly reduced in the mAb treated animals. In mAb 99 treated (1×10^5^ copies of N1/μg of RNA; p = 0.0264), P7 treated (4×10^4^ copies of N1/μg of RNA; p = 0.0188), Sotrovimab treated (9×10^3^ copies of N1/μg of RNA; p = 0.0170) and PBS only (2×10^6^ copies of N1/μg of RNA; Fig. 4**C**). A histological and immunohistological examination was performed on the lungs of all animals in this experiment. The PBS control group, revealed changes consistent with SARS-CoV-2 Delta infection in hamsters at this stage (*28*), represented by multifocal consolidated areas with evidence of epithelial regeneration (represented by hyperplastic bronchiolar epithelial cells), type II pneumocyte activation and hyperplasia, and a predominantly mononuclear inflammatory infiltrate that still contained a proportion of neutrophils (Fig. 5**A**). There was very limited viral antigen expression, in rare individual macrophages in consolidated areas, and in a few alveolar epithelial cells (Fig. 5A). In all treated hamster cohorts, there was no evidence of viral antigen expression in any of the lung sections. In the mAb 99 treated cohort, four animals exhibited either no or minimal non-specific changes (perivascular mononuclear leukocytes) (Fig. 5**B**), and there was no evidence that these animals had undergone a pulmonary infection. The remaining two animals (2.4, 2.5) as well as all the mAb P7 treated animals (Fig. 5**C**) exhibited small consolidated areas with epithelial hyperplasia similar to the PBS control cohort, consistent with prior focal SARS-CoV-2 infection that had induced epithelial cell damage and loss, followed by regenerative processes. In the Sotrovimab control animals, histological changes were only seen in 4/6 lungs (Fig. 5**D**); these were minimal and non-specific (perivascular mononuclear leukocytes). There was no evidence that this cohort of animals had undergone a pulmonary infection. Detailed information on individual animals is provided in Table **S2.**

**Figure 4.**
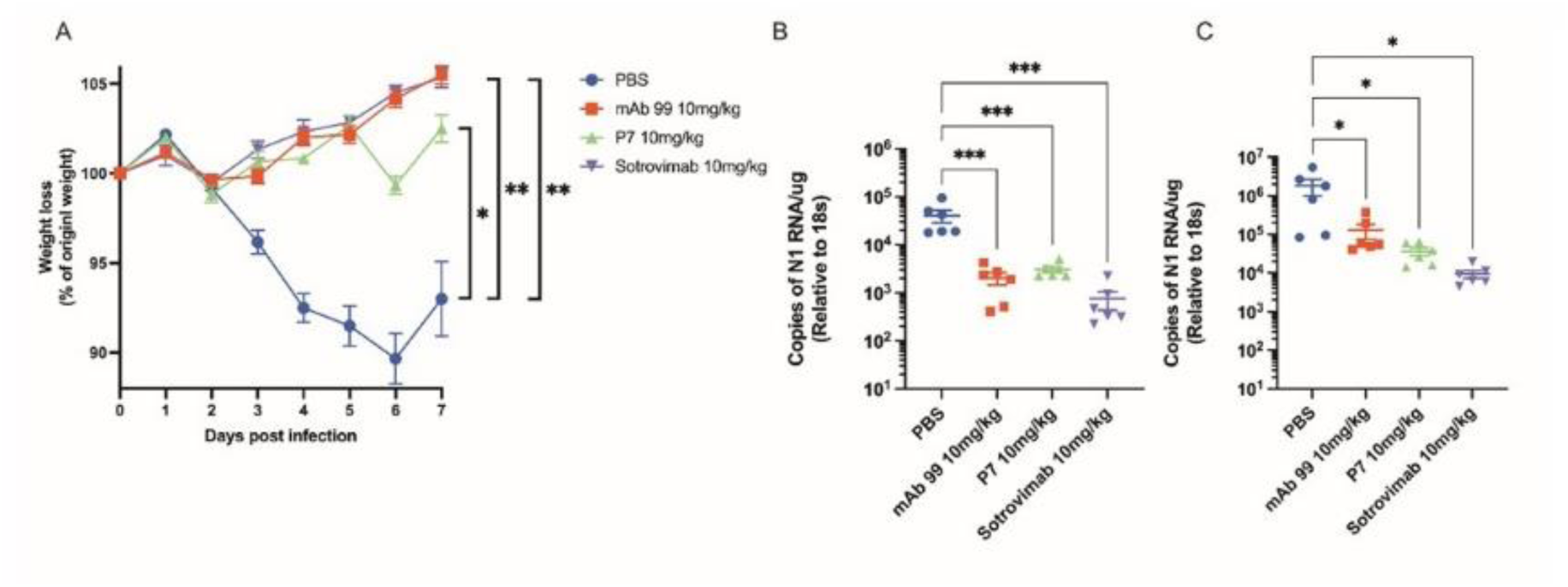
*In vivo* challenge in Syrian hamsters. (**A**) Golden Syrian hamsters (n = 6 biologically independent animals per group) were infected intranasally with SARS-CoV-2 strain Delta; 10^4^ pfu. Individual cohorts were treated 24 h post-infection (hpi) with 200 μL of antibody intranasally (IN) or sham-infected with PBS. Animals were monitored for weight loss at indicated time-points. Data are the mean value (n = 6) ± SEM. Comparisons were made using a repeated-measures two-way ANOVA with Geisser-Greenhouse’s correction and Dunnett’s multiple comparisons test; at day 7: PBS vs. mAb 99 10 mg/kg; **p = 0.0034, PBS vs. P7 10 mg/kg; ***p = 0.0109, PBS vs. 10 mg/kg Sotrovimab; **p = 0.0034 (**B**) RNA extracted from lungs was analysed for SARS-CoV-2 viral load using qRT-PCR for the N gene levels by qRT-PCR. Assays were normalised relative to levels of 18S RNA. Data for individual animals are shown with the mean value represented by a horizontal line. Data are mean value (n = 6) ±SEM and were analysed using a one-way ANOVA with Dunn’s multiple comparisons test; PBS vs. 10 mg/kg mAb 99; p = 0.0006, PBS vs. 10 mg/kg P7; p = 0.0008, PBS vs 10 mg/kg Sotrovimab; p = 0.0006. (**C**) RNA extracted from the nasal tissue was analysed for SARS-CoV-2 viral load using qRT-PCR for the N gene levels by qRT-PCR. Assays were normalised relative to levels of 18S RNA. Data for individual animals are shown with the mean value represented by a horizontal line. Data are mean value (n = 6) ±SEM and were analysed using a one-way ANOVA with Dunn’s multiple comparisons test; PBS vs. 10 mg/kg mAb 99; p = 0.0264, PBS vs. 10 mg/kg P7; p = 0. 0188, PBS vs 10 mg/kg Sotrovimab; p = 0. 0170

**Figure 5.**
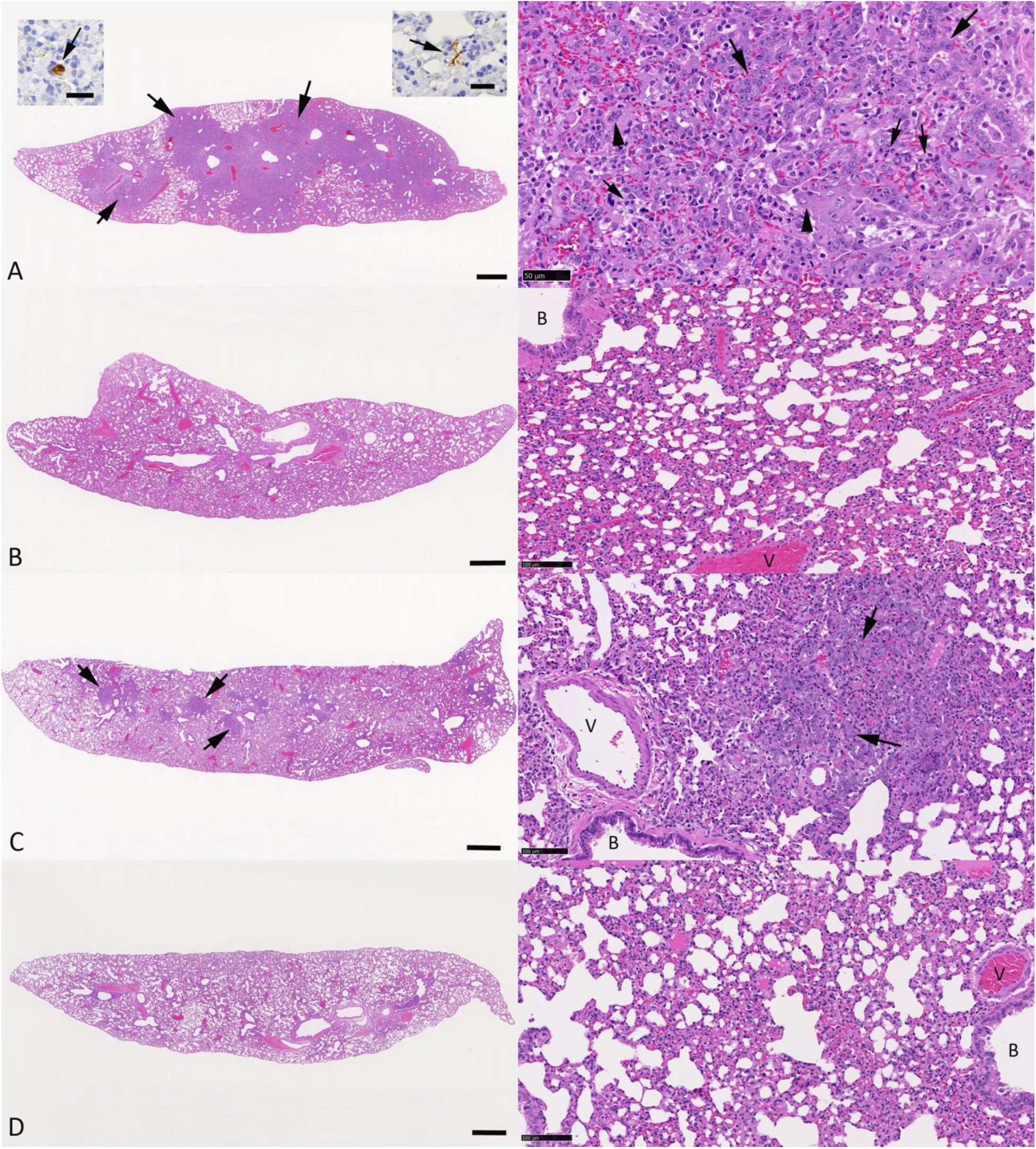
Histological changes in the lungs of male Syrian hamsters, aged 9-11 weeks, at day 7 post intranasal challenge with SARS-CoV-2 Delta at a dose of 10^4^ PFU that had received PBS (cohort 1), mAb 99 (cohort 2), mAB P7 (cohort 3) or Sotrovimab (cohort 4) 24 hours pre infection. Evidence of viral antigen expression is restricted to the animals from cohort 1. (**A**) Animal 1.5. The lung shows extensive multifocal consolidated areas (left image: arrows). The right image shows a higher magnification of a consolidated area, with hyperplastic bronchiolar epithelial cells (arrows), large activated type II pneumocytes (arrowheads) and embedded aggregates of neutrophils (small arrows). Viral antigen expression is restricted to very rare individual macrophages (left image: left inset, arrow) and rare alveoli with a few positive epithelial cells (left image: right inset, arrow). (**B**) Animal 2.6. The lung does not exhibit any histological changes. (**C**) Animal 3.6. The lung exhibits a few random focal consolidated areas (left: arrows). The right image shows a focal consolidated area with small aggregates of hyperplastic bronchiolar epithelial cells (arrows) and infiltrating leukocytes. (**D**) Animal 4.2. The lung does not exhibit any histological changes. HE stains. Bars = 1 mm (left column), 50 µm (right column: A) and 100 µm (right column: B-D). B: lumen of a bronchiole; V: lumen of a vein.

To further assess the *in vivo* efficacy of our antibodies, we evaluated mAb 99 and P7 at a lower dose (4 mg/kg) against live Omicron BA.5 virus in Syrian hamsters. Weight of hamsters remained stable during the experiment (data not shown) as previously reported (*29*). Measuring the lung viral load seven days post-infection showed that animals treated with mAb 99 (5×10^3^ copies of N1/μg of RNA; p = 0.0346) and P7 (3×10^3^ copies of N1/μg of RNA; p = 0.0096) had a significant decrease compared to the PBS only control (2×10^4^ copies of N1/μg of RNA). The control Sotrovimab was not significantly protective (7×10^3^ copies of N1/μg of RNA; p = 0.0856; fig. S7**A**).

We only tested mAb 99 against Lineage A *in vivo* as mAb P7 was isolated after the experiment was conducted. The animals in the untreated group showed a small but not significant progressive weight loss, whereas all animals treated therapeutically showed only a small weight loss 24 h after viral challenge, and from day 2 had recovered to at least pre-challenged weights (fig. S7**B**). The same result was observed in animals administered with the control commercial Ronapreve (casirivimab and imdevimab) mAb cocktail. In addition to measuring hamster weight loss, we measured viral loads in the lungs on day 7. Animals treated with mAb 99 exhibited a significantly lower viral load (3×10^3^ copies of N1/μg of RNA; p = 0.0007) than the PBS control (5×10^5^ copies of N1/μg of RNA). Ronapreve also had a significant reduction on viral load in the lungs (9×10^2^ copies of N1/μg of RNA; p = 0.0007; fig. S7**C**).

### Cryo-EM reveals insight into UL-CDRH3 binding

To further characterize the mechanism by which antibody 99 neutralises SARS-CoV-2, we solved a single-particle cryo-electron microscopy (cryo-EM) structure of the Fab from antibody 99 in complex with the SARS-CoV-2 Lineage A spike trimer at 3.6 Å resolution (Fig. 6**A**). In this structure, all three RBDs of the spike trimer were bound by 99 Fab, with two RBDs adopting “up” conformations and the third RBD adopting the “down” conformation. As RBDs in an up conformation are more flexible than those in the down conformation (*30, 31*), we locally refined the region corresponding to the 99 Fab bound to the down RBD to 4.1 Å resolution to resolve interactions between 99 Fab and the RBD (fig. **S8**). In addition, we determined a crystal structure of 99 Fab at 2.8 Å resolution to model the CDRs at higher resolution. As shown in Figure 6A, interactions between antibody 99 and the RBD are mediated by all CDR regions with the 28 residues non-UL CDRH3 contributing most of the interactions (Fig. 6A). Overall, antibody 99 recognizes the class 1 RBD epitope that overlaps with the ACE2 binding footprint (Fig. 6**B**, **C**) (*32*). Class 1 anti-RBD antibodies were previously defined as only targeting RBDs in up conformations (*32*); our finding of antibody 99 binding to a down RBD was possibly due to the flexibilities of the up RBDs that resulted in greater opening of the spike trimer, which allowed antibody 99 to bind to a down RBD. This is not the first time that a class 1 anti-RBD antibody was observed binding to a down RBD; for example, a potent human neutralising class 1 anti-RBD antibody, R40-1G8, was also seen to bind to the down conformation of the RBD (*33*).

**Figure 6.**
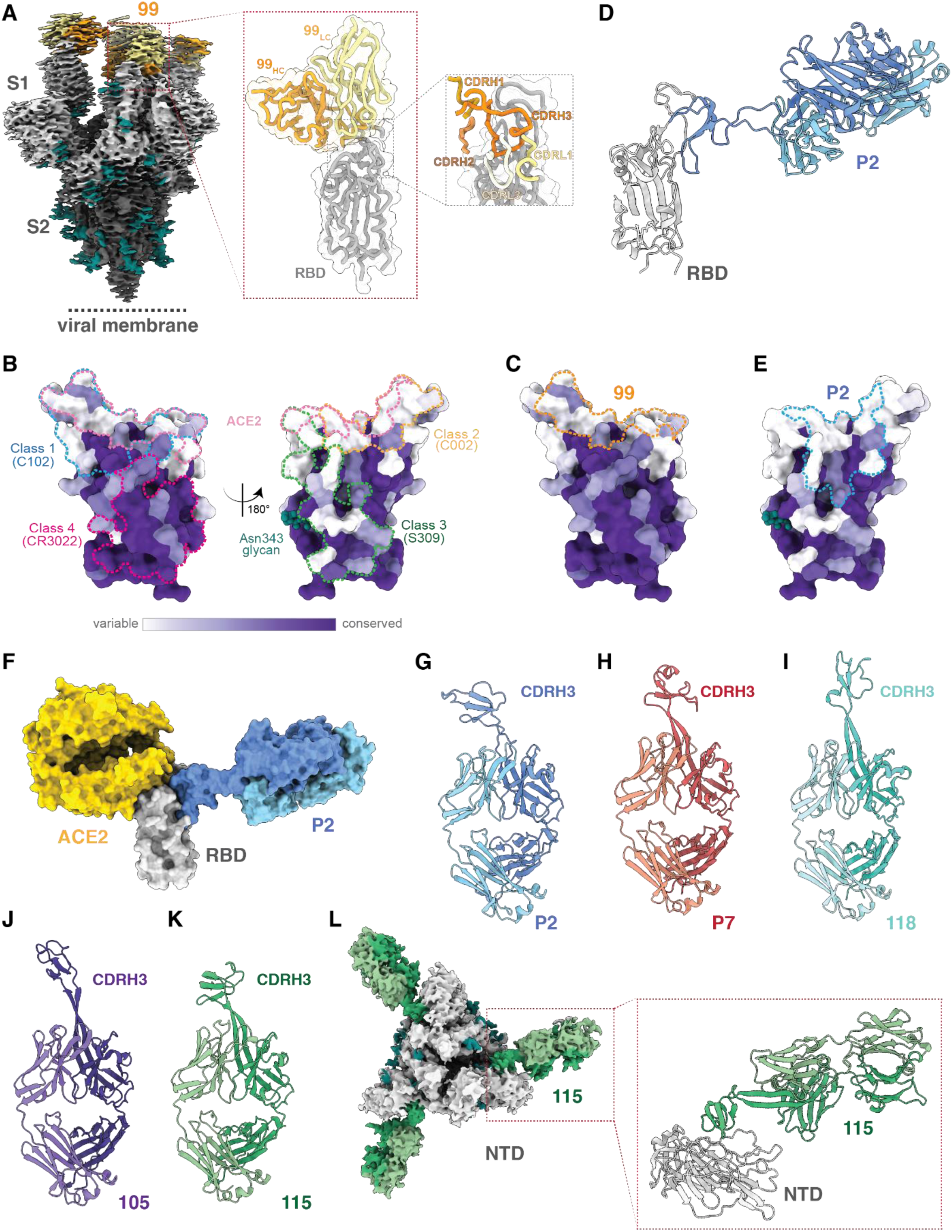
Interactions of bovine Fabs with SARS-CoV-2 spike. (**A**) Left EM density of single-particle cryo-EM structure of 99 Fab-spike trimer complex from side view. Details of 99 Fab-RBD complex (right). (**B**) Sequence conservation of 16 sarbecovirus RBDs calculated by ConSurf Database [59] plotted on an RBD structure (PDB 7BZ5). Epitopes for representative mAbs (C102, PDB 7K8M; C002, PDB 7K8T; S309, PDB 7JX3; CR3022, PDB 7LOP; and ACE2 PDB 6M0J are outlined in dotted lines. (**C**) Binding epitope of 99 Fab on the RBD sequence conservation plot. (**D**) 2.4 Å crystal structure of a P2 Fab-RBD complex. (**E**) Binding epitope of P2 Fab on the RBD sequence conservation plot. (F) Structural alignment of the ACE2-RBD (PDB 6M0J) and P2-RBD complexes. Crystal structures of (**G**) P2 from the P2-RBD crystal structure (2.4 Å resolution), (**H**) P7 Fab (1.7 Å resolution), (**I**) 118 Fab (1.9 Å resolution), (**J**) 105 Fab (1.65 Å resolution), and (**K**) 115 Fab (1.7 Å resolution). (**L**) Single-particle cryo-EM density of the 115 Fab-spike complex. Tsoleridis 2024 - An ultra-long heavy chain bovine antibody neutralizes SARS-CoV-2 and reacts broadly with sarbecoviruses.

We next investigated RBD recognition by the bovine antibodies with UL CDRH3s. We first solved a 2.4 Å resolution crystal structure of a complex between the P2 Fab and a SARS-CoV-2 Lineage A spike RBD (Fig. 6**D**). The RBD epitope of the P2 Fab (Fig. 6**E**) is adjacent to the epitope of the 99 Fab (Fig. 6C), which overlapped with the footprints of a class 2 anti-RBD mAb (C002) (*32*) and a class 3 mAb (S309) (*34*) footprint (Fig. 6B). There was no direct structural clash between ACE2 and P2 Fab on the RBD when we structurally aligned a complex structure with ACE2-RBD to the P2-RBD structure (Fig. 6**F**). We therefore classified P2 as a class 2 anti-RBD antibody as its binding footprint overlapped mostly with the class 2 mAbs. The interaction between P2 Fab and RBD is entirely mediated by its CDRH3, and the 63-residue UL CDRH3 of P2 folds into three antiparallel β-strands that are stabilized by disulfide bonds, and the strands are connected by two loops that interact with the RBD (Fig. 6**G**). This is in contrast with the RBD interactions mediated by the non-UL mAb 99 using all of its CDRs (Fig. 6A).

The conformation of the P2 CDRH3 (Fig. 6G) was different from the UL (61-residue) CDRH3 in the mAb P7, based on a 1.7 Å P7 Fab crystal structure (Fig. 6**H**). The P7 knob region folded into two antiparallel β-strands, connected by a long loop and one short helical turn (Fig. 6H). Although we were not able to obtain a structure of a P7 Fab-RBD complex, the interactions of P7 with the RBD are potentially mediated by the loops of the CDRH3 knob, in common with interactions of P2 and other bovine mAbs (*35*). The classification of the 99 and P2 antibodies as class 1 and class 2 anti-RBD antibodies, respectively, is consistent with the selective binding and neutralisation of SARS-CoV-2 VOCs for these antibodies (Fig. 1 and 2), since the class 1 and class 2 RBD epitopes exhibit more variability than other RBD epitopes, e.g., class 3 and class 4 RBD epitopes (Fig. 6B). On cryo-EM grids, P7 Fabs induced trimer dissociation, suggesting it binds to an RBD epitope that is not accessible in an intact spike trimer. Based on the cross-reactive RBD recognition of P7 by ELISA and neutralisation, P7 is likely a broadly cross-reactive antibody that targets the most conserved RBD region; i.e., the class 4 region that is buried on a down RBD. Further evidence for assigning P7 as a class 4 anti-RBD mAb is based on observations that class 3 RBD epitopes are generally more accessible in both RBDs adopting up and down conformations (*32*) and P7 does not compete for binding with mAb 99 or P2.

In addition to the anti-RBD mAbs with UL CDRH3s, we also solved crystal structures of three other Fabs from mAbs 105, 115, and 118 (Fig. 6**I**, **J**, **K**), all of which have UL CDRH3s. In common with P2 and P7 and other UL CDRH3 mAbs (*35*), the UL CDRH3s of these Fabs also folded into knob structures that are stabilized by disulfide bonds. We also solved a cryo-EM structure of a SARS-CoV-2 Lineage A spike trimer in complex with the 115 Fab (Fig. 6**L**). A low-resolution reconstruction revealed that mAb 115 targets the NTD near residues 23-28, 61-66, 80-87, and 136-139 (exact interactions were not resolved due to the relatively poor resolution of the EM map (fig. **S9**). Similar to the P2 Fab-RBD structure, the interaction between the 115 Fab and the NTD is solely mediated by its UL CDRH3.

## DISCUSSION

The constant evolution of SARS-CoV-2 within the human population gives rise to new VOCs that can more easily escape immune reactions in the current landscape of vaccinated and infected individuals. VOCs, such as Omicron, contain mutations in key residues of spike that affect the efficacy of currently available vaccines and neutralising antibodies (*18, 36*). Therefore, it is crucial to identify and develop broadly neutralising mAbs that confer protection against future variants of concern.

Here, we showed that consecutive immunisations with Lineage-A spike in cows, generated a strong and broad immune response against all SARS-CoV-2 variants, as well as against SARS-CoV-1. A recently published study confirms our findings of generation of broad potent bovine mAbs, showing that cow immunisations with the ancestral Lineage-A spike can elicit broad antibody responses, which neutralise contemporary SARS-CoV-2 variants and/or other sarbecoviruses (*35*). The strongest and most broad immune response was observed in the final timepoint D242, which was 73 days after the previous bleed on D169 (Fig. 1A and B), consistent with previous studies showing that longer intervals between later boosts induce strong and broad immune responses (*25, 37*).

We employed two different methods to retrieve SARS-CoV-2 spike-specific antibodies from bovine PBMCs: i) single spike-specific B-cell sorting in 96-well plates, and ii) bulk sorting of spike-specific B-cells and phage display. Both approaches have been used by previous studies to retrieve antibodies from cow PBMCs (*25, 38*). Our data showed that, within a panel of 33 antibodies, 21 bound to the SARS-CoV-2 spike, of which 5 recognised the RBD (Fig. 1C). The RBD-binding mAbs P2, P7 (both ULs), and 99 (non-UL) showed great breadth and binding capacity against SARS-CoV-2 VOCs. mAbs 99, 103, and 101 competed with human ACE2 for RBD binding, whereas P2 and P7 did not (Fig. 1D) suggesting neutralisation is achieved without complete obstruction of the ACE2 RBD interaction, similar to mAbs S2H97 and B9-scFv (*11, 26*). Individual RBD-targeting mAbs, such as Regdanvimab and Bamlanivimab (which compete with ACE2), and Sotrovimab, S2X259, and Bebtelovimab (which do not compete with ACE2) Licensed therapeutic mAbs targeting RBD, such as Regdanvimab and Bamlanivimab, are ACE2-competitive mAbs, while Sotrovimab, S2X259, and Bebtelovimab are not (*34, 39–41*), as well as mAb cocktails such as Evusheld (cilgavimab + tixagevimab) and Ronapreve (casirivimab + imdevimab) have been used as therapeutics for SARS-CoV-2 (*42, 43*), although their lack of effectiveness against some VOCs has led to their removal from clinical use. This highlights the need to isolate alternative, next-generation mAbs to overcome the threat of escape.

Further characterisation of the 21 spike-binding mAbs in single-point neutralisation assays against pseudotyped SARS-CoV-2 spikes revealed the best and most broadly neutralising mAbs. mAbs P2, P7, 118, 105 (ULs) and 99 (non-UL) were selected and further characterised because they could neutralise at least 5 out of 6 variants, with a minimum of 75% efficacy (Fig. 2A). Comparison of the *in vitro* IC_50_ values of our mAbs to the licenced therapeutic mAbs showed that mAbs P2 and 99 had similar neutralisation potencies to Amubarvimab, Romlusevimab, and Sotrovimab for variants Alpha, Beta, and Delta (*42*). Licenced antibodies Sotrovimab and Romlusevimab showed a 3-fold reduced potency against the Omicron (B.1.1.529) variant, whereas most of the other therapeutic antibodies except for Bebtelovimab, showed great reductions in neutralisation against this variant (*42, 44*). In contrast, our antibodies were still in the range of IC_50_ 10-50 ng/mL against Omicron. The mAb P2 did not neutralise BA.5 suggesting mutations within the RBD for this subvariant were key residues in the binding epitope of this mAb. Similar class 2 mAbs, such as bamlanivimab (LY-CoV555), were also reported to display substantial evasion by BA.5 (*17, 45*). The mAbs P7 and 99 were able to potently neutralise more contemporary variants such as Omicron BA.5 and XBB (Fig. 2B and C). The most broadly neutralising mAb was P7, which neutralised SARS-CoV-1 and bound RBDs across several *sarbecovirus* clades, suggesting it targets a conserved epitope within sarbecovirus RBDs (Fig. 3). Recently published studies have characterised similar SARS-CoV cross-neutralising mAbs such as S2X259, S2K146, Bebtelovimab and E7 (*12, 41, 46, 47*).

In addition to SARS-CoV-1 cross-neutralisation, mAb P7 also showed cross-reactivity against RBD present in representative animal sarbecoviruses drawn from clades 1a, 1b, 2 and-3. Pan-coronavirus cross-recognition has been observed for some monoclonals targeting the NTD of spike, but the reported affinity and potency of these antibodies is low (*48, 49*). Several broadly reactive pan-sarbecovirus RBD-targetting murine and human mAbs have been reported, and these include S309 (Sotrovimab), 2-36, M2-7, M8a-31, M8a-34, CC25.54 and CC25.36 (*14, 34, 50–52*).

A common feature of these human RBD-targeting mAbs with broad reactivity is their relatively extended CDRH3 (>20 amino acids). Given P7’s ultralong CDRH3, it is tempting to speculate that possession of a CDRH3 with increased length gives favourable cross-reactivity, possibly by enabling engagement with highly conserved epitopes that are less accessible to mAbs with shorter CDRH3s. Unfortunately, we were not able to resolve the structure of mAb P7 in association with spike as the Fab resulted in trimer dissociation.

The efficacy of the candidate mAbs *in vivo*, showed a level of protection that was at least equivalent to commercial mAb therapeutics; Ronapreve (a cocktail of casirivimab and imdevimab, both IgG1) against Lineage A and Sotrovimab against Delta and Omicron BA.5 (Fig. 4). Of note is that the antibodies presented here do not have Fc modifications to improve their in vivo half-life unlike Sotrovimab (LS mutation (Met428Leu/Asn434Ser)) (*53*), suggesting that they may be even more efficacious with the same modifications.

Our data showed that, although the cows were immunised with Lineage-A spike, we discovered antibodies that could bind several animal *sarbecovirus* RBDs. Assessment of the binding capability of our RBD-binding mAbs against a panel of *sarbecovirus* RBDs showed that the most broadly reactive mAb was P7 (23 of 23), followed by 99 (8 of 23), P2 (7 of 23), 103 (4 of 23) and 101 (3 of 23) (Fig. 2). In a recent study, scientists discovered broadly neutralising antibodies able to bind multiple *sarbecovirus* RBDs from the same panel as ours, by immunising mice with mosaic RBD nanoparticles (*14*). Another recently published study reported the isolation of potent pan huACE2-dependent sarbecovirus neutralising mAbs from a BNT162b2-vaccinated SARS-CoV-1 survivor (*54*). Although mAb 99 exhibited strong neutralisation potencies across different SARS-CoV-2 VOCs, it was not as broadly reactive as P7, which was identified as a pan-*sarbecovirus* antibody. Overall, our study not only demonstrates the isolation of neutralising mAbs from immunised cows, but also highlights the discovery of two new broadly neutralising mAbs. These antibodies show great potential for identifying conserved epitopes, and as therapeutic candidates against SARS-CoV-2, with P7, in particular, standing out as a pan-*sarbecovirus*-reactive antibody that could serve to inform vaccine design, as a diagnostic tool, and as a pandemic response therapeutic.

## MATERIALS AND METHODS

### Immunisations

Two Holstein bullocks were immunised each with 250 µg of the S1 subunit of SARS-CoV-2 spike (REC31806, Native Antigen Company) mixed with Titremax Gold Adjuvant (Sigma Aldrich). A booster immunisation was performed with 150 µg of the same protein/adjuvant combination 21 and 42 days after the prime immunisation. Another booster immunisation was performed 161 days post-prime, with 250 µg of SARS-CoV-2 spike ectodomain (S1 and partial S2; aa 1-1211, SRAS furin cleavage site, transmembrane domain and intravirion part replaced with a glycine-serine linker; REC31868, Native Antigen Company) mixed with Titremax Gold Adjuvant (Sigma Aldrich). A final boost was performed on day 235 with 200 µg of SARS-CoV-2 trimeric spike (aa 1-1208, GSAS furin cleavage site, soluble 2P (aa 986-987), purity and size verified by manufacturer; REC31871, Native Antigen Company) mixed with the same adjuvant. Blood (50 mL) was collected on days 25, 46, 50, 165, 169, 239 and 242 – approximately 4-and 8-days post immunisation. Also, blood was collected on day 0 as a control.

### Blood processing

Peripheral blood mononuclear cells (PBMC) and plasma were harvested after processing the blood samples with Histopaque-1077 (Sigma-Aldrich) and SepMate PBMC isolation tubes (Stemcell Technologies). PBMCs and plasma were stored in LN_2_ and-80 °C, respectively. The plasma samples were heat inactivated at 56 °C for 40 min before use in assays.

### Flow cytometry sorting

Twenty million (20 x 10^6^) PBMCs were defrosted for each bullock and incubated O/N at 37 °C in 5% CO_2_ in Gibco™ RPMI 1640 Medium supplemented with 10% heat inactivated fetal bovine serum (FBS). The cells were washed with 3% bovine serum albumin (BSA, Sigma) dissolved in phosphate-buffered saline (PBS) with the addition of 2 mM ethylenediaminetetraacetic acid (EDTA, Sigma). After centrifugating at 300g for 10 minutes and then washing again, the cells were stained with biotinylated Lineage A trimeric spike (The Native Antigen Company) for 30 min at 4 °C followed by washing, centrifugating and staining with R-PE and APC conjugated streptavidin (Thermo Fisher Scientific) for 30 min at 4 °C. After the final wash, the cells were stained with Annexin-V FITC and propidium iodide (PI, Thermo Fisher Scientific) for 20 min at 4 °C. Using an Astrios cell sorter (Beckman Coulter), cells were gated using FSC/SSC plots and aggregates excluded by SSC pulse width. The lymphocyte gate was identified based on forward and side scatter. Doublets were then excluded, and Annexin-V FITC and PI cell dyes were used for live-dead discrimination. R-PE^+^ and APC^+^ spike-specific labelled B-cells were gated as our primary population of interest and were sorted into 96-well plates containing 1 mg/mL BSA (Thermo Fisher Scientific) and were frozen immediately at-80 °C. In addition to single cells, a bulk sort of live spike-specific B-cells was also performed for each bullock, which was then used for phage display.

### Retrieval of antibody heavy and light chains and cloning

After the sorted single cells were snap frozen, cDNA was synthesised using RNA to cDNA EcoDry Premix Random Hexamers (Takara Bio) following the manufacturer’s instructions. Full-nested PCR was performed to retrieve the heavy and light chain for each B-cell using HotStartTaq DNA polymerase (Qiagen) following the manufacturer’s conditions. The first round was a multiplex targeting the IgG heavy chains and the lambda light chains (IgL). The second round PCRs were targeting IgG and IgL separately. Each IgH PCR product was cloned into an IgG1 cassette in pcDNA3.1, which was comprised of human constant region and bovine variable regions. Similarly, the IgL PCR products were cloned into a human IgL cassette containing bovine variable regions. The cloning was performed using In-Fusion cloning (Takara Bio) following the manufacturer’s protocol. The Fab fragment versions of the cloned antibodies were generating by removing their Fc region using a Q5 Site Directed Mutagenesis kit (New England Biolabs).

### Generation of Phage Libraries from bulk sorted anti-specific B cells

RNA was extracted from bulk sorted SARS-CoV-2 spike reactive B-cells using an RNA extraction kit (Qiagen), according to manufacturer’s instructions. Purified RNA was used as the template for cDNA synthesis using EcoDry Premix random hexamers (Takara Bio). Heavy chain variable regions of VH1-7 family were amplified from the cDNA by PCR using VH family specific primers. Retrieved VH1-7 amplicons were subcloned into a linearised phagemid pFab-VL vector by Gibson assembly (New England BioLabs) according to manufacturer’s instructions. The pFAb-VL vector encoded a bovine CL and CH1 framework and a previously described VL1-47 light chain variable region (*25*). The bovine Fab was placed upstream of a truncated phage gene 3 sequence to allow incorporation of Fab into phage particles on the p3 minor protein. Products of the Gibson assembly were electroporated in TG1 *E. coli* cells (Lucigen) and transformed cells were grown on 2XYT supplemented with 2% glucose and 100 µg/mL ampicillin (2XYT GA) agar plates and incubated at 37 °C overnight.

Phagemid libraries were rescued by harvesting colonies from each culture plates into 25 mL 2XYT GA media and grown until log phase when M13K07 helper phage (New England BioLabs) was added at an MOI 1.25. Cultures were incubated with helper phage for 45 min at 37 °C and then pelleted by centrifugation at 3000 *x* g for 10 min. Supernatant was discard and pelleted cells were resuspended in glucose free media and grown for 18 h at 30 °C. Phage particles were harvested from infected cultures by removal of bacterial cells by centrifugation followed addition of PEG precipitation buffer (20 % PEG8000, 2.5 M NaCl). Culture supernatants were incubated in PEG buffer overnight at 4 °C and precipitates were pelleted by centrifugation at 4000 *x* g for 10 min followed by resuspension in PBS. Phage particles were quantified by infecting TG1 *E. coli* followed by titration and inoculation of 2XYT GA agar plates to calculate colony forming units per µL.

### Phage Biopanning

Phage libraries were normalised to a concentration of 1 x10^11^ cfu/well and added to a 96 well maxisorp plate (Thermo Fisher Scientific) that had been coated with trimeric SARS-CoV-2 spike from either Lineage A (first round) or BA5 (second round) variants, at a concentration of 5 µg/mL and blocked with 3% BSA for the first round, and 3% Gelatin from cold water fish skin. Phage particles were incubated on the plate for 1 hr at room temperature with gentle agitation followed by 20 washes with PBST with increasing concentrations of Tween20 from 0.1% to 1.0%. Bound phage particles were eluted from the plate by the addition of acidic glycine buffer (0.2 M glycine, p*H* 2.2) and incubated for 10 minutes followed by neutralisation with 1 M Trizma hydrochloride (Tris HCL, p*H* 9.1). Eluted phage was added to 600 µL of TG1 cells that were at log-phase and cells were cultured for 1 hour then plated on 2XYT GA agar. Colonies were phage rescued and titrated 24 hrs later as previously described.

### Mammalian cell culture

HEK-293T and Vero-E6 adherent cell lines were maintained in DMEM supplemented with 10 % FBS (Sigma-Aldrich) and 1 % non-essential amino acids (Gibco) and grown at 37 °C with 5% CO_2_. The HEK derived suspension cell line, Expi293 (Thermo-Scientific) was maintained in Expi293 cell media (Thermo-Scientific) and propagated at 37 °C, 8% CO_2_ with agitation at 125 rpm.

### Expression of recombinant mAbs

Small scale expression of the mAb panel was initially performed using transient transfection of HEK-293T cells. In brief, 4 µg of heavy chain-encoding plasmid and 4 µg of light chain-encoding plasmid were prepared for transfection by mixing with 16 µg P3000 reagent (Thermo Fisher Scientific), 24 µg Lipofectamine reagent (Thermo Fisher Scientific) and Opti-MEM (Gibco), made up to a final volume of 500 µL. This solution was incubated for 20 min at room temperature and was added to a 10 cm Primaria™ coated dishes (Corning) that had been seeded with 3×10^6^ HEK-293T cells 24 hrs prior to transfection. The cells were incubated for 6 hrs and the media was replaced with 10 mL of fresh DMEM. Transfected cultures were grown for 72 hrs and the supernatant was harvested and passed through a 0.22 µM syringe filter (Sartorious).

For larger scale recombinant mAb production, the Expi293 expression system (Thermo Fisher Scientific) was used. Expi293 cells cultures were seeded at 2.5 x10^6^ cell/mL 24 hrs prior to transfection and incubated at 37^°^C, 8% CO_2_ with shaking at 125 rpm. Cell densities were adjusted to 3 x10^6^ cells/mL and transfected with heavy and light chain encoding plasmids at a ratio of 1:2, respectively, with the total amount of plasmid equivalent to a final concentration of 1 µg/mL of DNA in the transfected culture. Cultures were harvested 7 days post-transfection and centrifuged at 300 *x* g for 20 min to remove cells followed by centrifugation at 4500 *x* g for 1 hr to remove subcellular debris.

### Purification of recombinant mAbs

Supernatants harvested from mAb expressing Expi-293 cultures were diluted at a 1:1 ratio with binding buffer (100 mM sodium phosphate, p*H* 7.0) and passed through a 0.22 µm filter (StarLab). Recombinant mAb was purified from samples using an AKTA start system (Cytiva) equipped with a 1 mL HiTrap™ Protein G HP column (Cytiva). All steps were performed at a flow rate of 2 mL/min. In brief, the protein G column was equilibrated with 10 column volumes (CVs) prior to addition of the sample. Sample was applied to the column followed by a washing step with 15 CVs of binding buffer. Captured mAb was eluted from the column using 15 CVs of elution buffer (200 mM glycine, p*H* 2.7). Eluted sample was collected as 1 mL fractions that were mixed with 200 µL of neutralisation buffer (1 M Tris, p*H* 9.0). Fractions that contained the elution peak were pooled and concentrated using an Ultra-4 centrifugal filter device (Amicon) with a molecular weight cut-off of 100 KDa. Removal of residual glycine was achieved by buffer exchange using a zeba column (Thermo fisher scientific) equilibrated with PBS. Purified mAbs were then stored at-20 °C for future use.

### IgG ELISA quantification assay

Recombinant mAbs were quantified using an in-house IgG ELISA. In brief, 96 well maxisorp ELISA plates (Thermo Fisher Scientific) were coated with anti-human IgG heavy chain antibody (I2136-1ML, Sigma-Aldrich) at a concentration of 10 µg/mL, diluted in carbonate-bicarbonate buffer, at a volume of 50 µL/well. Following overnight incubation at 4 °C, coating buffer was removed, and plates were blocked with Phosphate-Buffered Saline with 0.1% Tween 20 (PBST) and 5% skimmed milk for 2 hrs at room temperature then washed once with PBST buffer. Samples were serially diluted, alongside a standard of recombinant human IgG1 Lambda antibody (Sigma-Aldrich) that was used to generate a standard curve starting from 20 µg/mL and diluted 2-fold for 16 points. Standards and samples were incubated on the plate for 90 min at room temperature and were removed followed by three washes with PBST. Captured IgG was detected by a 1 hr incubation with an anti-human Lambda chain Alkaline phosphatase secondary antibody at a 1:2000 dilution in PBST. Following removal of the secondary antibody, ELISA plates were washed a further three times with PBST and detection of bound antibody complexes were measured by the addition of para-Nitrophenylphosphate (pNPP) (Sigma-Aldrich) colorimetric substrate with absorbance read at 405 nM. IgG concentration of samples was interpolated from the standard curve.

### Expression of *sarbecovirus* RBD panel

A panel of *sarbecovirus* RBDs similar to those described in (*14*) were expressed in HEK 293T cells. Briefly, 4 µg of plasmid encoding a His-tagged RBD was mixed with p3000 and lipofectamine 3000 (Thermo Fisher Scientific), according to the manufacturer’s instructions, and transfected into a 10 cm dish of 3 x 10^6^ HEK 293T cells. Supernatant was harvested 72 hrs post transfection and soluble RBD proteins were purified using nickel-affinity purification. All centrifugation steps were carried out at 1000 *x* g for 1 minute. In brief, supernatant was incubated with 200 µL of Ni-NTA His bind resin (Merck) and incubated for 1 hr at room temperature. A resin solution was added to a 10 KDa Amicon Ultra-0.5 device (Merck) and supernatant removed by centrifugation. Resin was washed in 50 mM phosphate buffer (p*H* 8.0, 10 mM Imidazole) and flow through was discarded. The sample was added to the exchange device and incubated for 1 hour at room temperature with gentle agitation. Unbound protein was removed by centrifugation and flow through discarded. The device was subsequently washed with wash buffer, centrifuged and flow through discarded. Bound protein was eluted from the resin by addition of elution buffer (50 mM phosphate buffer, p*H* 8.0, 0.3NaCl, 25 mM Imidazole) and incubated for 5 minutes at room temperature. The eluted protein was collected by centrifugation at 4000 x g for 15 minutes. Eluted protein was validated by SDS-PAGE and silver staining.

### Binding ELISA

SARS-CoV-2 antigens were coated onto 96 well maxisorp plates (Thermo-Scientific) at a concentration of 500 ng/mL, and 5 µg/ml for the panel of 23 *sarbecovirus* RBDs, and incubated at 4 °C overnight. Plates were blocked with 5% skimmed milk PBST for 2 hrs. Recombinant antibodies were serially diluted 3-fold with a starting concentration of 2.5 µg/mL in duplicate and incubated on the ELISA plate for 1 hr. The wells were washed three times with PBST and anti-human IgG-Fc specific alkaline phosphatase (A3187, Sigma) was added at a 1:2000 dilution followed by 1 hr incubation. The assay was developed using SIGMAFAST^TM^ p-Nitrophenyl phosphate (pNPP) Tablets (Sigma) according to manufacturer’s procedure and absorbance read at an optical density of 405 nm, using a Fluorostar Omega microplate reader (BMG Labtech, Aylesbury, UK), in 5-minute intervals until saturation was reached. Analyses of results were carried out using GraphPad Prism version 10 for macOS (Graphpad Software, San Diego, USA).

Analyses as proportion of binding (%) relative to the maximum binding (2.5 μg/mL antibody concentration) and minimum binding negative control)). The EC_50_ was calculated using a non-linear analysis model.

ACE2 inhibition ELISA was performed similar to the binding ELISA, except recombinant antibodies were serially diluted four-fold in ACE2/PBST starting at a concentration of 2.5 μg/mL in duplicate and incubated on the ELISA plate for 1 hr. The wells were washed three times with PBST and Monoclonal Anti-Goat/sheep Alkaline Phosphatase antibody produced in mouse (Sigma: A8062) was added at a 1:2000 dilution followed by 1 hour incubation. The assay was developed and analysed as previously mentioned.

### Retroviral pseudotype particle neutralisation assay

Neutralising activities of bovine sera and recombinant monoclonal antibodies were assessed using a SARS-CoV-2 retroviral pseudotype (SARS-CoV-2 pp) assay, as described previously (*55*). In brief, 1.5 x10^6^ HEK-293T cells were seeded in a 10 cm Primaria coated dish (Corning) in 10 mL of DMEM, 24 hrs prior to transfections. The following day, 2 µg of MLV *gag-pol* packaging plasmid (phCMV-5349), 2 µg MLV-luciferase plasmid (pTG126) and 2 µg of pcDNA3.1 plasmid (Invitrogen) encoding SARS-CoV-2 spike proteins, were mixed with 24 µL polyethylenimine (Polysciences) diluted in Opti-MEM (Gibco) and incubated for 1 hr a room temperature. Plasmids were added to cells and incubated for 6 hrs at 37 °C and then removed and replaced with 10 mL of DMEM. Supernatants of transfected cell cultures were harvested 72 hrs post-transfection and passed through a 0.45 µM syringe filter (Sartorius) and stored at 4 °C and used within 1 week of harvesting. For neutralisation assays, 270 µL of SARS-CoV-2 pps were mixed with either 30 µL of diluted recombinant antibody or diluted heat inactivated Bovine plasma and incubated at room temperature for 1 hr. SARS-COV-2 pps were then added in triplicate to a white 96 well plate (Corning) that had been seeded with 2 x10^4^ Vero-E6 cells 24 hrs prior. Vero-E6 cells were incubated with SARS-CoV-2 pps for 4 hrs at 37 °C followed by removal of pseudotyped viruses and addition of 200 µL of DMEM. Vero-E6 cells were incubated for a further 72 hrs at 37°C, 5 % CO_2_. Luciferase activity was measured using the Promega luciferase assay system, cell supernatant was removed from the 96 well plate and 50 µL of cell lysis buffer was added per well and incubated a room temperature for 30 min with agitation followed by addition of 50 µL luciferase substrate solution per well. Luminescence was measured using a FLUOstar Omega plate reader (BMG Labtech). Analyses of results were carried out using GraphPad Prism version 10 for macOS (Graphpad Software, San Diego, USA). Luminescence was normalised to positive and negative controls, calculated as proportion of neutralisation relative to the maximum infectivity (positive control) and minimum infectivity (delta Envelope pseudovirus particles). The IC_50_ for curves was calculated on GraphPad Prism by transforming the data and using a non-linear analysis model.

### Authentic virus neutralisation assay

Authentic SARS-CoV-2 neutralisation assays were performed as previously described (*56*) except that 3.95 x10^3^ TCID_50_ of virus was added to each well and each antibody was diluted from 5 µg/mL down a 4-fold dilution series in triplicate. For this assay, antibodies P7 and 99 were tested against SARS-CoV-2 variants Lineage A, Alpha, Beta, Delta and Omicron.

### *In vivo* infection studies

Animal work was approved by the local University of Liverpool Animal Welfare and Ethical Review Body and performed under UK Home Office Project Licence PP4715265. Male golden Syrian hamsters (*Mesocricetus auratus*) aged between 8 and 10 weeks old, weighing 90-100 g were purchased from Janvier Labs (France). Animals were maintained under SPF barrier conditions in individually ventilated cages. For virus infections the following strains were used: Liverpool strain. a PANGO lineage B strain of SARS-CoV-2 (hCoV-2/human/Liverpool/REMRQ0001/2020) was cultured from a nasopharyngeal swab from a patient, was passaged in Vero E6 cells (Patterson et al., 2020). The B.1.617.2 (Delta variant) hCoV-19/England/SHEF-10E8F3B/2021 (GISAID accession number EPI_ISL_1731019) and BA.5 was provided by Prof. Wendy Barclay, Imperial College London, London, UK through the Genotype-to-Phenotype National Virology Consortium (G2P-UK). All virus stocks had been checked for deletions in the mapped reads and the stocks confirmed to not contain any deletions.

Animals were randomly assigned into multiple cohorts of 6. For SARS-CoV-2 infection, hamsters were anaesthetised lightly with 5% isoflurane and inoculated intranasally with 100 µl containing 10^4^ PFU of SARS-CoV-2 Delta variant and 10^5^ PFU of Liverpool and BA.5 variants in PBS. At day-1 pre-challenge (pc), for the Delta variant infection, hamsters were treated with 10 mg/kg of mAbs 99 and P7 via the intraperitoneal route (I/P) in 200 µl PBS. Control hamsters (n = 6) received either no treatment (negative control) or Sotrovimab at 10 mg/kg body weight as a positive control. For BA.5, hamsters were treated at day-1 pc with 4 mg/kg of mAbs 99 and P7 via the intraperitoneal route (I/P) in 200 µl PBS. Control hamsters (n = 6) received either no treatment (negative control) or Sotrovimab at 4mg/kg body weight as a positive control. For the Liverpool variant, hamsters were treated at day-1 pc with 4 mg/kg of mAb 99 via the intraperitoneal route (I/P) in 200 µl PBS. Control hamsters (n = 6) received either no treatment (negative control) or Ronapreve (REGN-COV2; 10 mg/kg body weight I/P) as a positive control.

Hamsters were monitored daily for weight loss. Oropharyngeal (throat) swabs were obtained on days 3 and 5 post infection. Animals were briefly anaesthetised with 5% isoflurane for the collection of these samples. On day 7, all hamsters were euthanised by an overdose of anaesthetic (sodium pentobarbitone) via the intraperitoneal route. At necropsy, nasal turbinates and tissue samples (right lung) were collected and stored frozen at −80 °C for viral RNA measurement. From animals that had been infected with SARS-CoV-2 Delta and were treated with mAb 99, mAb P7 and Sotrovimab, the left lung was fixed in 10% buffered formalin for histological and immunohistological examination.

### RNA Extraction and qRT-PCR

RNA was isolated from oropharyngeal swabs and tissue samples (lung and nasal turbinates). Right upper lung lobe and nasal turbinates were homogenised in 1 ml of Trizol reagent (Thermofisher) using a QT tissue lyser and stainless-steel beads (Qiagen) at 50 oscillation for 5 minutes whereas swabs samples were inactivated in 1:3 ratio of Trizol LS (Thermofisher). The homogenates were clarified by centrifugation at 12,000xg for 5 min before full RNA extraction was carried out according to manufacturer’s instructions. RNA was quantified and quality assessed using a Nanodrop (Thermofisher) before a total of 1µg was DNase treated using the TURBO DNA-free™ Kit (Thermofisher) as per manufacturer’s instructions.

Viral load was quantified using a 1 step RT-qPCR kit (Promega). For quantification of SARS-COV-2 the nCOV_N1 primer/probe mix from the SARS-CoV-2 (2019-nCoV) CDC qPCR Probe Assay (IDT) were utilised and murine 18s was used as a housekeeping gene to normalise the qPCR. The 18s standard was generated by the amplification of a fragment of the murine 18S cDNA using the primers F: ACCTGGTTGATCCTGCCAGGTAGC and R: AGC CAT TCG CAG TTT TGT

AC. Likewise, SARS-COV2 genomic N1 standard was generated by PCR using the qPCR primers. cDNA was generated using 1 step qRT-PCR kit (Promega) as per manufacturer’s instructions. Both PCR products were purified using the QIAquick PCR Purification Kit (Qiagen) and serially diluted 10-fold from 10^10^ to 10^4^ copies/reaction to form the standard curve.

### Histological and immunohistological examination

After fixation in 10% buffered formalin for 48 h, the left lungs of all hamsters were transferred into 70% ethanol until further processing. The lung was halved by a longitudinal section and routinely paraffin wax embedded. Consecutive sections (3-4 µm) were prepared and stained with hematoxylin eosin (HE) for histological examination or subjected to immunohistological staining. Immunohistology was performed to detect SARS-CoV-2 antigen, using the horseradish peroxidase (HRP) method and the following primary antibody: rabbit anti-SARS-CoV nucleocapsid protein (Rockland, 200-402-A50) as previously described (*57*).

### Protein expression and purification

The HexaPro (6P) version of SARS-CoV-2 Lineage A spike trimer (*58*), SARS-CoV-2 Lineage A RBD, Fabs from mAbs P2, P7, 105, 115, and 118, and 99 IgG were expressed in Expi293 cells by transient transfection as described (*32*). Proteins were purified following previously-established protocols (*14*). Briefly, His-tagged spike trimer was purified from Expi293 cell supernatant using a Ni-NTA column followed by size exclusion chromatography (SEC) using a Superose 6 10/300 column (Cytiva). Peak fractions were collected and concentrated to ∼6 mg/mL and flash frozen in 30 µL aliquots and stored in-80°C until use. His-tagged SARS-CoV-2 RBD and P2, P7, 10,5 115 and 118 Fabs were purified from cell supernatants using a Ni-NTA column, and eluents from NiNTA purifications were subjected to SEC using a Superdex 200 10/300 Increase column (Cytiva). For making RBD-Fab complexes, both proteins were co-expressed during a transient transfection in Expi293 cells. Peak fractions from SEC were pooled and concentrated to ∼25 mg/ml and screened for crystallization. 99 IgG was purified by a MabSelect SuRe column followed by SEC purification using a Superdex 200 10/300 Increase column (Cytiva). 99 Fab was generated by papain digestion from purified 99 IgG as described (*33*).

### Single-particle cryo-EM sample preparation, data collection, and structure determinations

SARS-CoV-2 Lineage A spike and 99 Fab were incubated at room temperature for 30 minutes at a spike protomer:Fab molar ratio of 1:1.1 with the final concentration of spike at 2 mg/mL. Fluorinated octylmaltoside (Anatrace) at a final concentration of 0.02% (w/v) was immediately added to the complex before freezing. Three µL of complex-detergent mixture was added to freshly glow-discharged QuantiFoil 300 mesh 1.2/1.3 grids (Electron Microscopy Sciences). Grids were blotted with zero blot force for three seconds at 100% humidity and room temperature using Whatman No.1 filter paper and vitrified in 100% liquid ethane using a Mark IV Vitrobot (Thermo Fisher Scientific).

Single-particle cryo-EM datasets for 99 Fab bound to the SARS-CoV-2 Lineage A spike 6P, and 115 Fab bound to the SARS-CoV-2 Lineage A spike 6P were collected on a 300 keV Titan Krios (Thermo Fisher Scientific) equipped with a K3 camera (Gatan) using SerialEM automated data collection software (*59*). Movies were recorded with a total dosage of 60 e^-^/Å^2^ in 40 frames using a defocus range of-1 to-3 µm and a pixel size of 0.416 Å in the super-resolution mode via a 3×3 beam image shift pattern with three exposures per hole. A detailed workflow for data processing of the spike-99 Fab complex and the spike-115 Fab are outlined in Supplementary Figure 8 and 9, respectively. Patch motion correction for movies was done using a binning factor of two, and contrast transfer function (CTF) parameters were estimated with patch CTF in cryoSPARC v3.3.25 (*60*). Particles were picked with blob picker using a particle diameter of 100-200 Å and extracted upon inspection in cryoSPARC (*60*). Particles were initially classified with 2D classification, and after discarding ice and junk particles, the remaining particles were used for ab initio model building with four volumes in cryoSPARC (*60*). These volumes were then used for heterogeneous refinements. After discarding junk particles and particles with preferred orientations, homogenous and non-uniform refinements were carried out for a final reconstruction in cryoSPARC (*60*). A mask of a Fab-RBD complex was generated using Chimera (*61*) and used for local refinement in cryoSPARC (*60*).

An initial model of the Fab-RBD region of the 99 Fab-spike complex structure was generated by docking the V_H_ of the bovine antibody BOV-3 (PDB 6E9H), the V_L_ of bovine antibody 2G3 (PDB 8ECQ), and the RBD of a previously-determined spike structure (PDB 7UZ6) into a locally refined cryo-EM map using UCSF Chimera (*61*). The model was refined into the local cryo-EM reconstruction using Coot (*62*) and real space refinement in Phenix (*63*). The refined 99 Fab-RBD and a previously-determined spike structure (PDB 7UZ6) were docked into the full single-particle cryo-EM map. Iterative real space refinement and model building were carried out in Phenix (*63*) and Coot (*62*), and the 99 Fab sequence was manually corrected using Coot (*62*). Refinement statistics are reported in Supplementary Table 3. Structure figures were made using UCSF ChimeraX (*64*).

### X-ray crystallography

Crystallization trials for unbound Fabs and Fab-RBD complexes were set up using commercially available screens by mixing 0.2 µL well solution with 0.2 µL of a protein solution at 10 mg/mL or 20 mg/mL using a TTP LabTech Mosquito instrument at room temperature. Crystals for RBD-P2 were obtained from a PEG Rx screen (Hampton Research) containing 0.1 M citric acid pH 3.5, and 25% w/v polyethylene glycol 3,350. Crystals for P7 Fab were obtained from a PEG ion screen (Hampton Research) containing 0.2 M sodium bromide and 20% w/v polyethylene glycol 3,350. Crystals for 99 Fab were obtained from a JCSG screen (Molecular Dimensions) containing 0.1 M HEPES 7.0 and 30 % v/v Jeffamine® ED-2003. Crystals for 105 Fab were obtained from a Morpheus screen (Molecular Dimensions) containing 0.12 M alcohol, 0.1M buffer system 3 pH 8.5, and 30% precipitant mix 1. Crystals for 115 Fab were obtained from Crystal screen (Hampton Research) containing 0.1 M HEPES sodium pH 7.5, 10% v/v 2-propanol, and 20% w/v polyethylene glycol 4,000. Lastly, crystals for 118 Fab were obtained from PEGion (Hampton Research) containing 0.1 M HEPES sodium pH 7.0, 0.01 M magnesium chloride hexahydrate, 0.005 M nickel chloride hexahydrate, and 15% w/v polyethylene glycol 3,350. All crystals were cryo-protected in well solution mixed with 20% glycerol before cryo-preservation in liquid nitrogen.

X-ray diffraction data were collected at the Stanford Synchrotron Radiation Lightsource beamline 12-2 and Advanced Light Source beamline 2.0.1. All X-ray datasets were indexed and integrated with XDS (*65*), and scaled with Aimless (*66*). The structure of the P2-RBD complex was solved by molecular replacement using structures of an RBD (PDB 7UZC) and a cow antibody (PDB 6E9H) as input models for *Phaser* in Phenix (*67*). The refined structure of the P2 Fab in the RBD-Fab complex was further used as a molecular replacement model for the the P7, 105, 115, and 118 Fab datasets. For the crystal structure of the 99 Fab, the refined V_H_-V_L_ from the single-particle 99 Fab-spike cryo-EM structure and the C_H_-C_L_ from the P2 Fab structure were used as input models for molecular replacement. For all crystal structures, iterative refinement and model-building cycles were carried out with phenix.refine in Phenix (*67*) and *Coot* (*68*), respectively. Crystallographic data collection and refinement statistics are reported in table **S4**.

## List of Supplementary Materials

Present a list of the Supplementary Materials in the following format.

Materials and Methods

Fig. S1 to S10

Tables S1 to S4

## Acknowledgments

We thank Jost Vielmetter, Luisa Segovia, Annie Rorick, and the Caltech Beckman Institute Protein Expression Center for protein production, Songye Chen and the Caltech Cryo-EM facility for cryo-EM data collection, and Jens Kaiser, staff at the Stanford Synchrotron Radiation Lightsource and Advanced Light Source, and the Caltech Molecular Observatory for X-ray data collection support. Cryo-Electron microscopy was performed in the Beckman Institute Resource Center for Transmission Electron Microscopy at Caltech. We also thank Dr David Onion and the University of Nottingham Flow Cytometry Facility. The contents of this publication are solely the responsibility of the authors and do not necessarily represent the official views of NIGMS or NIH.

## Funding

Use of the Stanford Synchrotron Radiation Lightsource, SLAC National Accelerator Laboratory, is supported by the U.S. Department of Energy, Office of Science, Office of Basic Energy Sciences under Contract No. DE-AC02-76SF00515.

The SSRL Structural Molecular Biology Program is supported by the DOE Office of Biological and Environmental Research, and by the National Institutes of Health, National Institute of General Medical Sciences (P30GM133894).

Beamline 2.0.1 of the Advanced Light Source, a DOE Office of Science User Facility under Contract No. DE-AC02-05CH11231, is supported in part by the ALS-ENABLE program funded by the National Institutes of Health, National Institute of General Medical Sciences, grant P30 GM124169-01.

National Institutes of Health P01-AI138938-S1 (PJB) National Institutes of Health P01-AI165075 (PJB)

Medical Research Council MR/S009434/1 and MR/R010307/1 (JKB) Biotechnology and Biological Sciences Research Council BB/M018636/1 (JKB)

## Author contributions

Conceptualization: TT, CF, AIF, PJB, RAU, JKB

Methodology: TT, EJP, CF, JDD, CMC, AIF, CPM, DXJ, MAAP, DH, RAU, JKB

Formal analysis: TT, EJP, CF, JDD, RAU, JKB

Investigation: TT, EJP, CF, JDD, PS, SW, AK, EGB, AK, XH, CMC, CPM, SR, JPS, PJB, RAU, JKB

Resources: IS, DB, DXJ, MAAP Data curation: TT, CF, JDD, RAU

Visualization: TT, CF, JDD, JGC, PJB, RAU Funding acquisition: TT, CPM, JPS, PJB, RAU, JKB

Project administration: TT, JDD, JPS, PJB, RAU, JKB Supervision: TT, JDD, AWT, JPS, PJB, RAU, JKB

Writing – original draft: TT, CF, JDD, PJB, RAU, JKB

Writing – review & editing: TT, EJP, CF, JDD, PS, SW, JGC, AK, EGB, AK, CMC, AIF, CPM, DXJ, MAAP, AWT, DH, JPS, PJB, RAU, JKB

## Competing interests

Authors declare that they currently have no competing interests. Patent filing in progress.

## Data and materials availability

All data are available in the main text or the supplementary materials.

## Supplementary methods

None

## Supplementary Figures

**Supplementary Figure 1.**
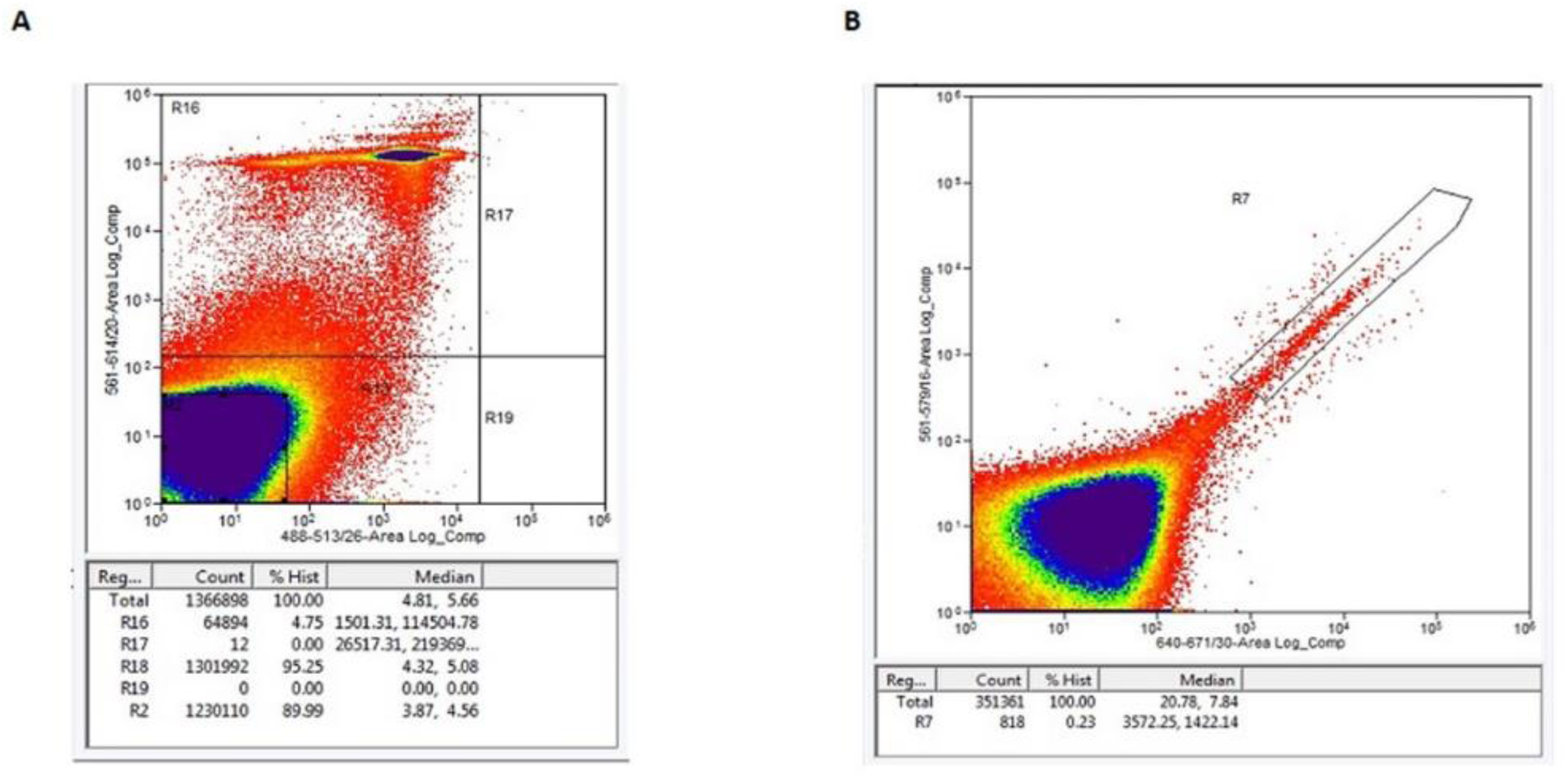
Gating strategy for PBMCs subjected to flow cytometry. (A) Viability gating with Annexin-V-FITC on the x-axis and Propidium Iodide on the y-axis. Cells in gate R2 were considered live and used for downstream sorting. (B) Sorting gate for SARS-CoV-2 spike double-positive live cells. X-axis represents APC stain and y-axis represents PE stain. Cells positive for both stains (gate R7) were sorted into a 96-well plate as single cells.

**Supplementary Figure 2.**
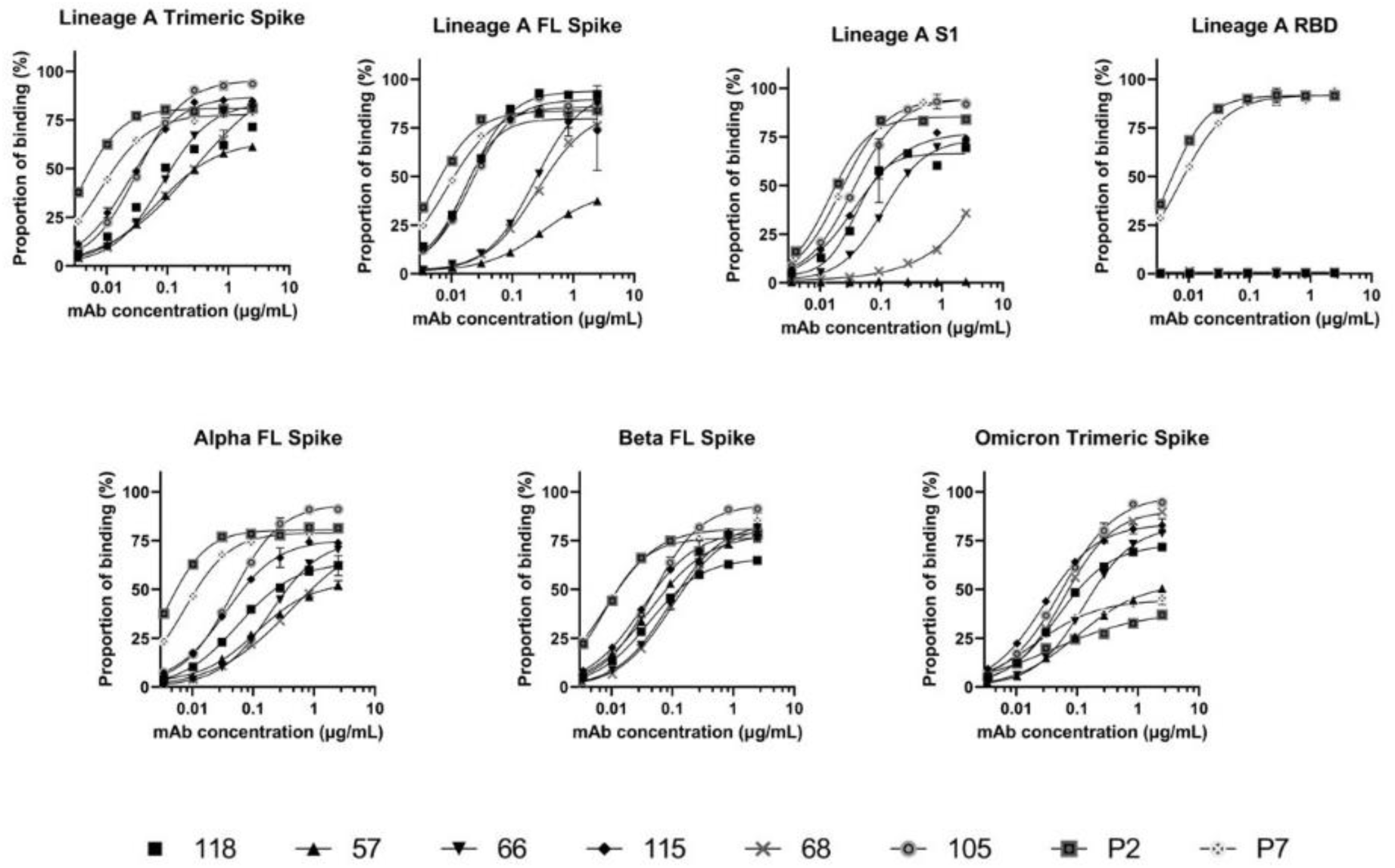
Binding curves of the eight ultra-long mAbs of the panel against different SARS-CoV-2 spike variants and forms of spike (Lineage A: trimer, full-length, S1, RBD, Alpha full-length, Beta full-length, Omicron trimer). The highest concentration of each mAb was 2.5µg/mL and the lowest was 3ng/mL

**Supplementary Figure 3.**
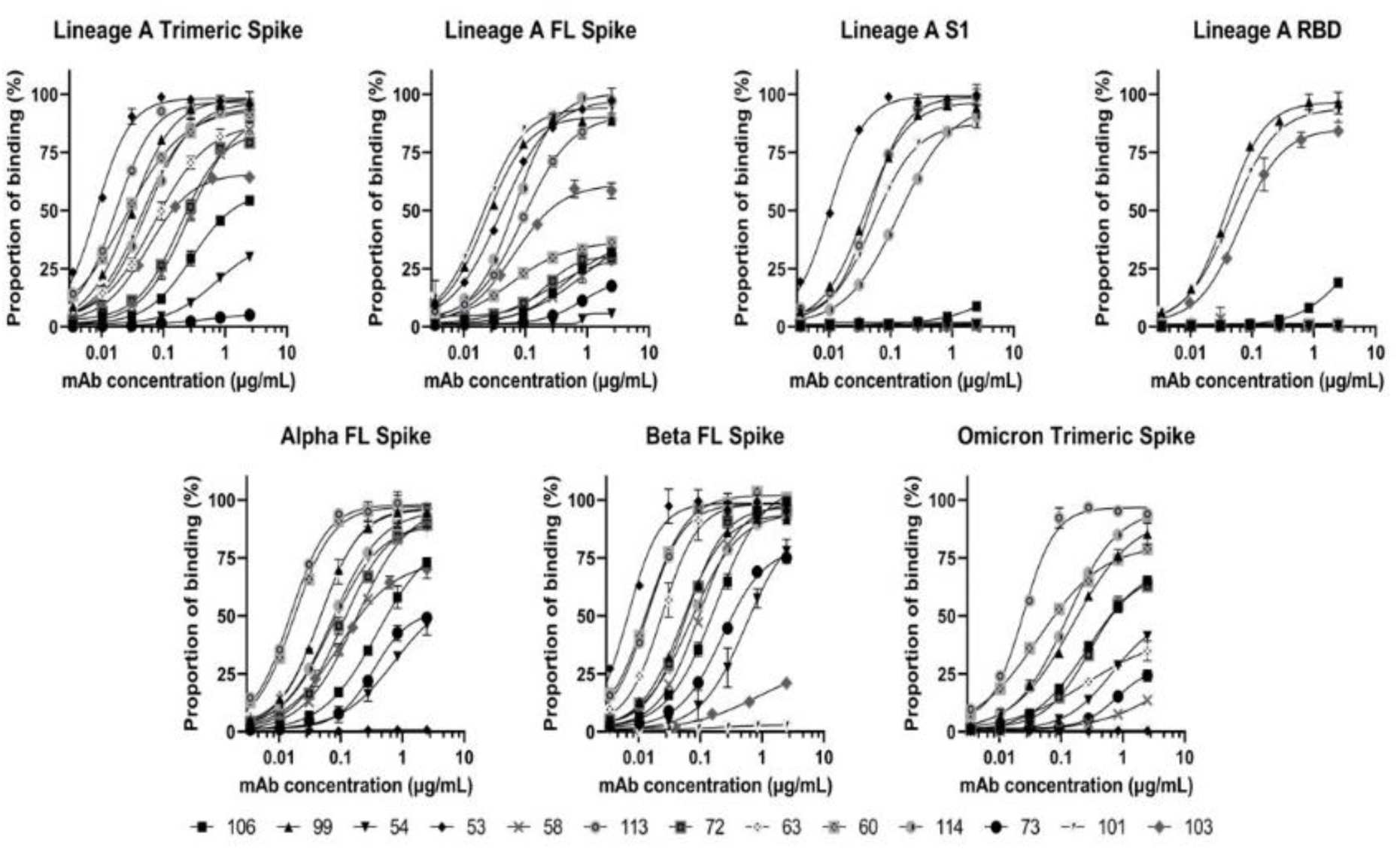
Binding curves of the 13 non-ultra-long mAbs of the panel against different SARS-CoV-2 spike variants and forms of spike (Lineage A: trimer, full-length, S1, RBD, Alpha full-length, Beta full-length, Omicron trimer). The highest concentration of each mAb was 2.5µg/mL and the lowest was 3ng/mL.

**Supplementary Figure 4.**
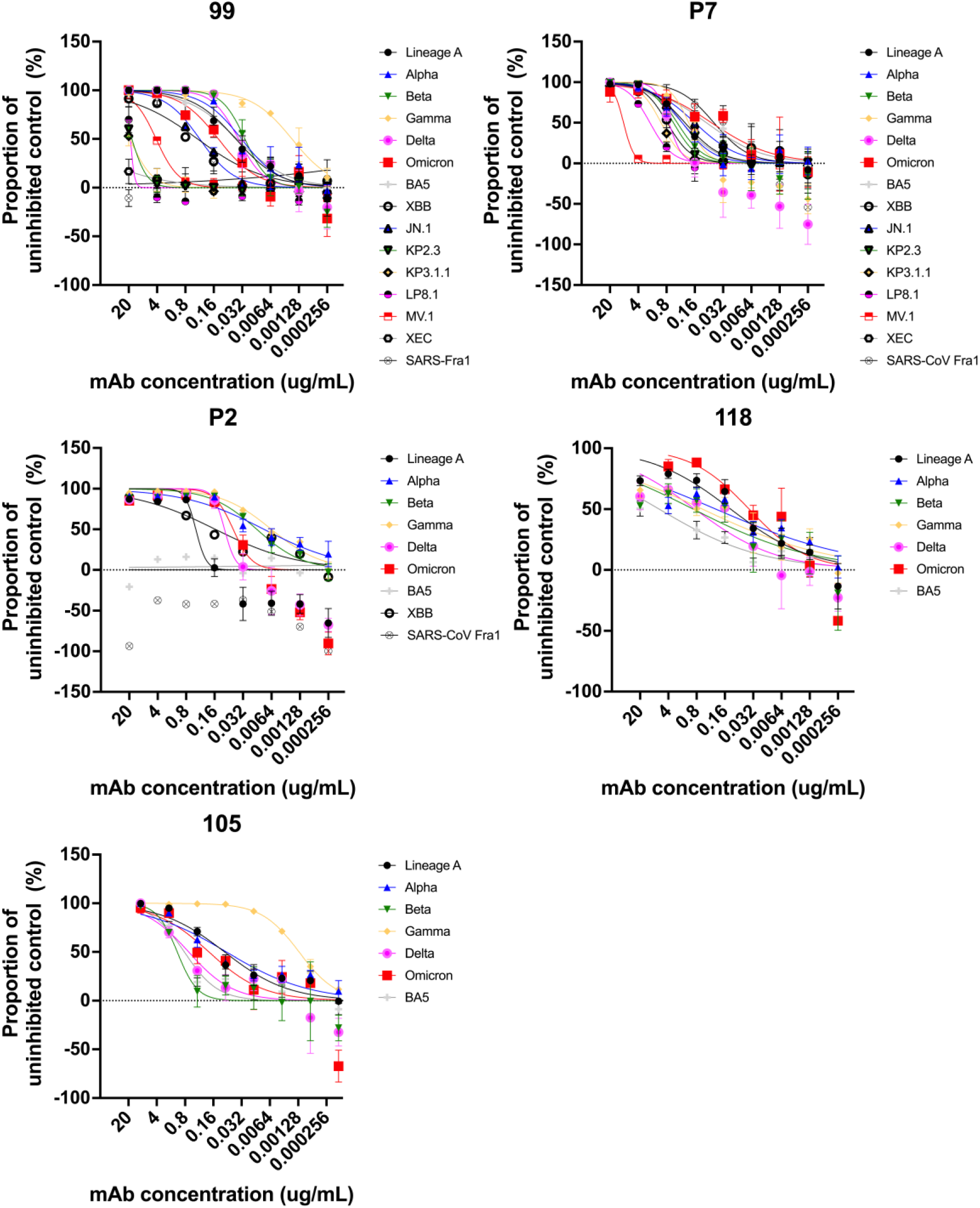
Neutralisation curves of antibodies 99 (non-UL), P7, P2, 118 and 105 (UL) against pseudotypes of SARS-CoV-2 variants.

**Supplementary Figure 5.**
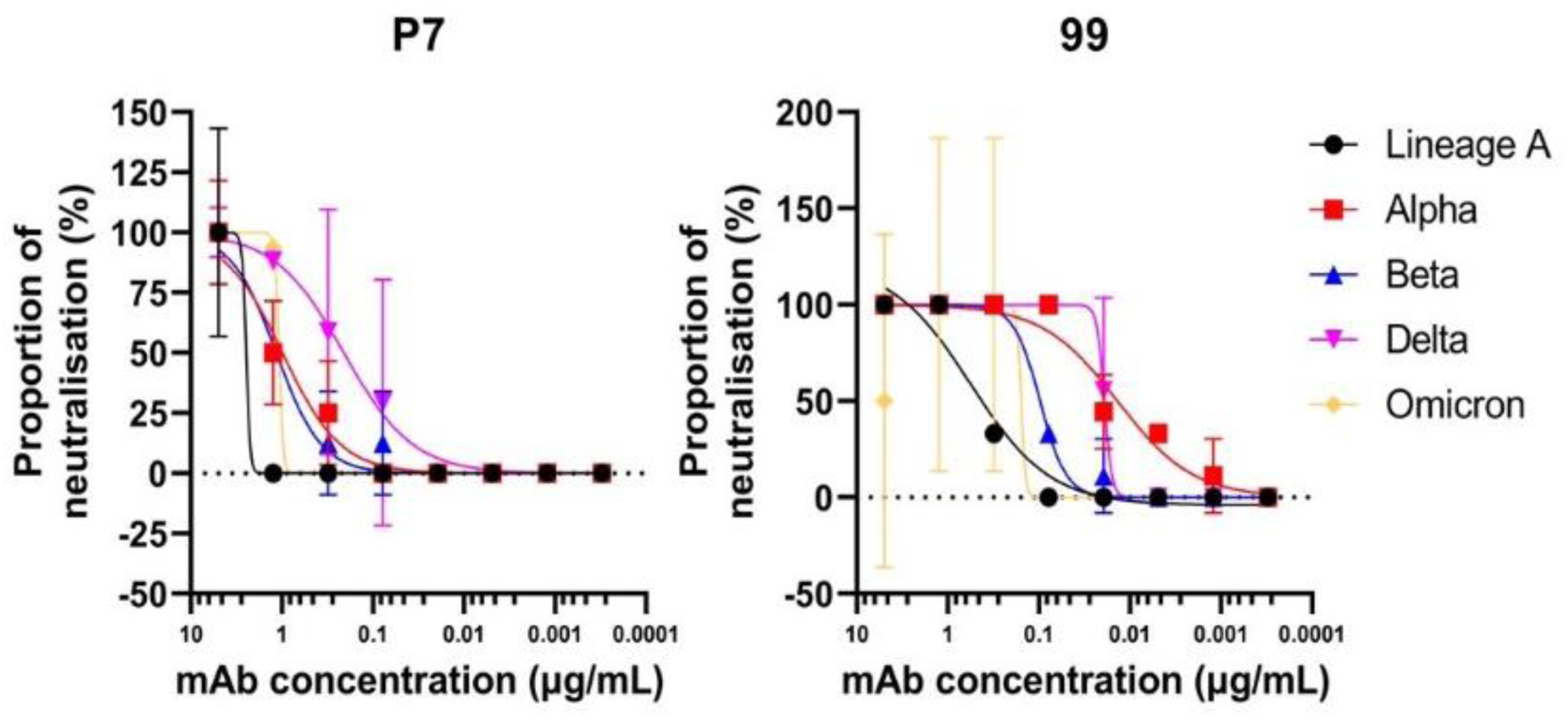
Neutralisation curves of mAbs P7 (UL) and 99 (non-UL) against different variants of authentic SARS-CoV-2.

**Supplementary Figure 6.**
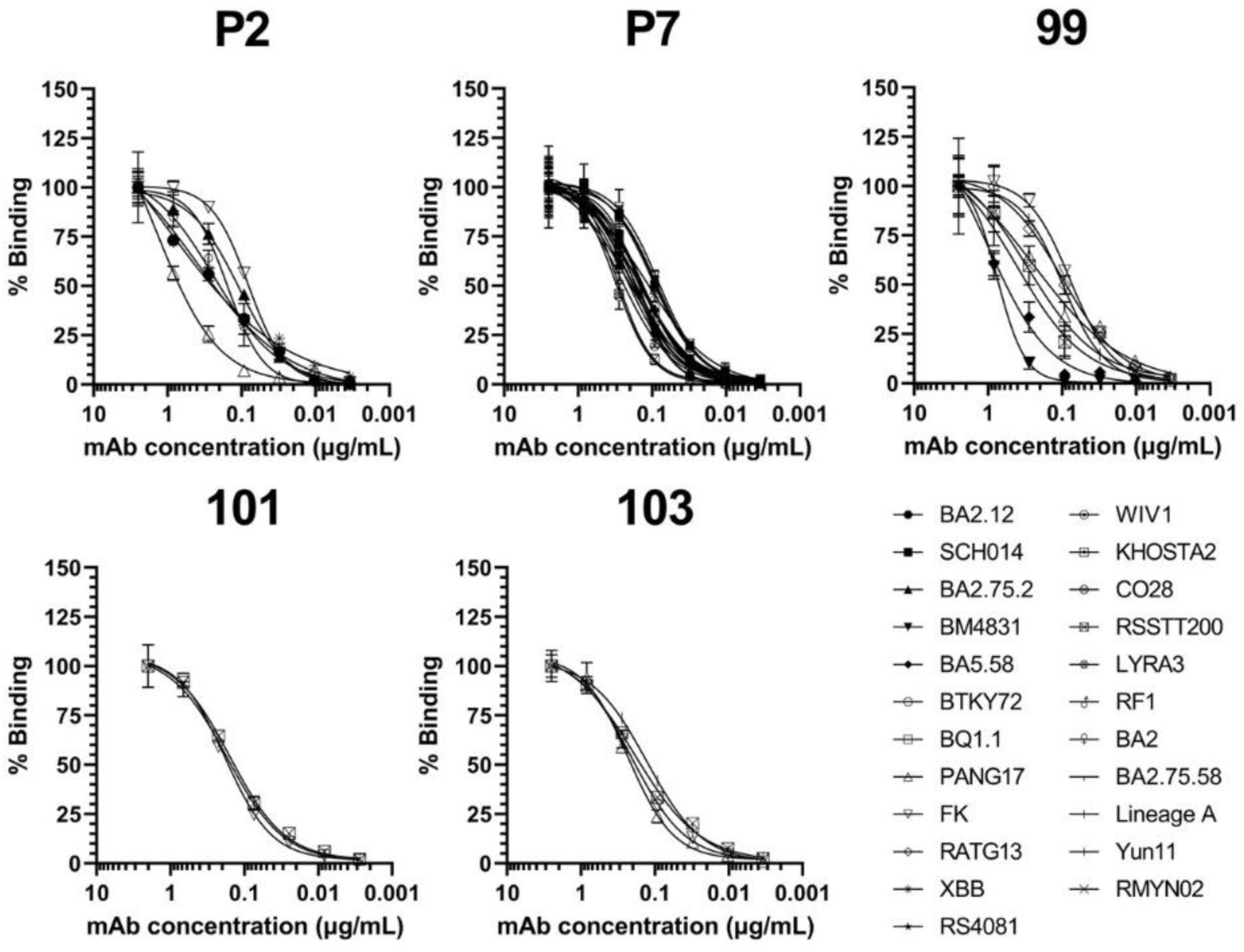
Binding curves of the RBD-binding mAbs against the RBD panel. Graphs show only the variants with detectable binding to the mAbs.

**Supplementary Figure 7.**
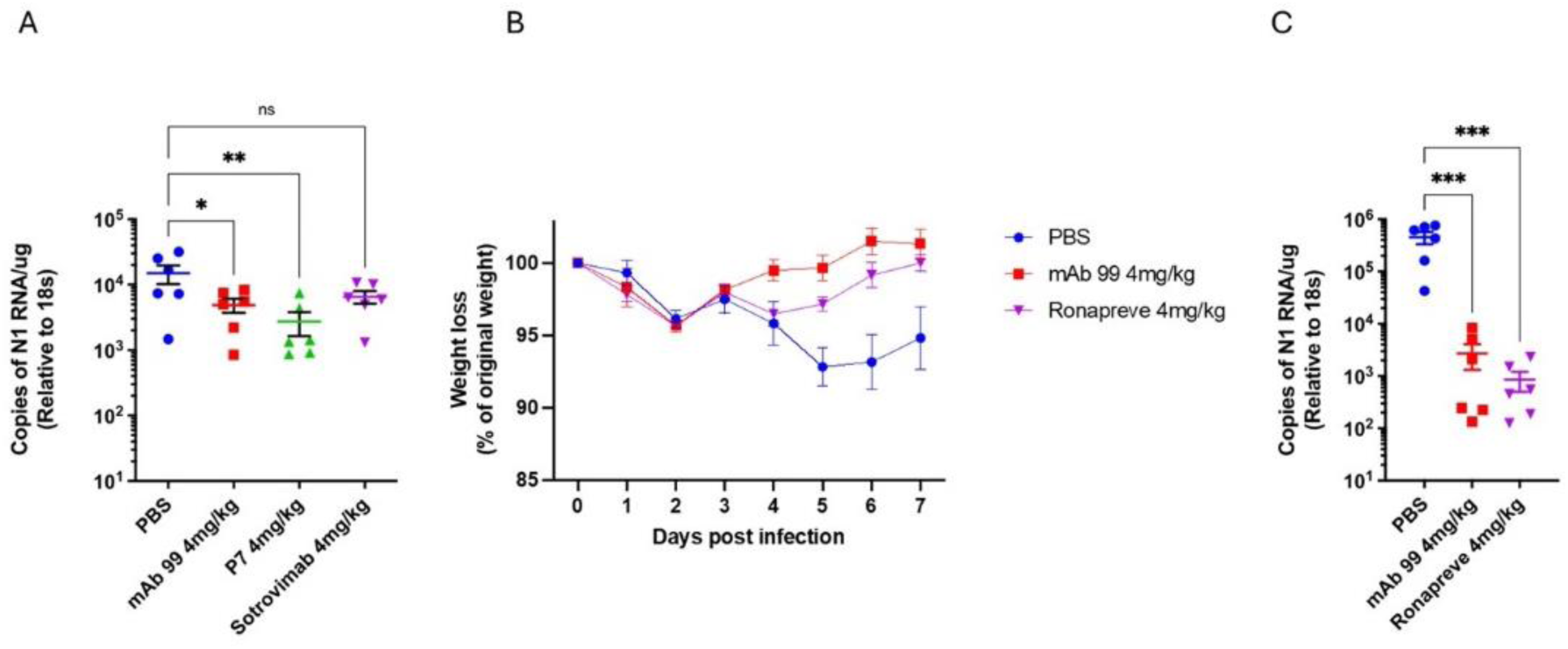
(A) Golden Syrian hamsters (n = 6 biologically independent animals per group) were infected intranasally with SARS-CoV-2 strain BA.5; 10^4^ pfu. Individual cohorts were treated 24 h post-infection (hpi) with 200 μl of antibody intranasally (IN) or sham-infected with PBS. RNA extracted from lungs was analysed for SARS-CoV-2 viral load using qRT-PCR for the N gene levels by qRT-PCR. Assays were normalised relative to levels of 18S RNA. Data for individual animals are shown with the mean value represented by a horizontal line. Data are mean value (n = 6) ±SEM and were analysed using a one-way ANOVA with Dunn’s multiple comparisons test; PBS vs. 4 mg/kg mAb 99; *p = 0. 0346, PBS vs. 4 mg/kg P7; **p = 0. 0096, PBS vs 4 mg/kg Sotrovimab; ns p = 0. 0856. (B) Golden Syrian hamsters (n = 6 biologically independent animals per group) were infected intranasally with SARS-CoV-2 strain LIVERPOOL; 10^4^ pfu. Individual cohorts were treated 24 h post-infection (hpi) with 200 μL of antibody intranasally (IN) or sham-infected with PBS. Animals were monitored for weight loss at indicated time-points. Data are the mean value (n = 6) ± SEM. Comparisons were made using a repeated-measures two-way ANOVA with Geisser-Greenhouse’s correction and Dunnett’s multiple comparisons test; at day 7 there were no significant differences. (C) RNA extracted from the lung tissue was analysed for SARS-CoV-2 viral load using qRT-PCR for the N gene levels by qRT-PCR. Assays were normalised relative to levels of 18S RNA. Data for individual animals are shown with the mean value represented by a horizontal line. Data are mean value (n = 6) ±SEM and were analysed using a one-way ANOVA with Dunn’s multiple comparisons test; PBS vs. 4 mg/kg mAb 99; p = 0. 0007, PBS vs 4 mg/kg Sotrovimab; p = 0. 0007.

**Supplementary Figure 8.**
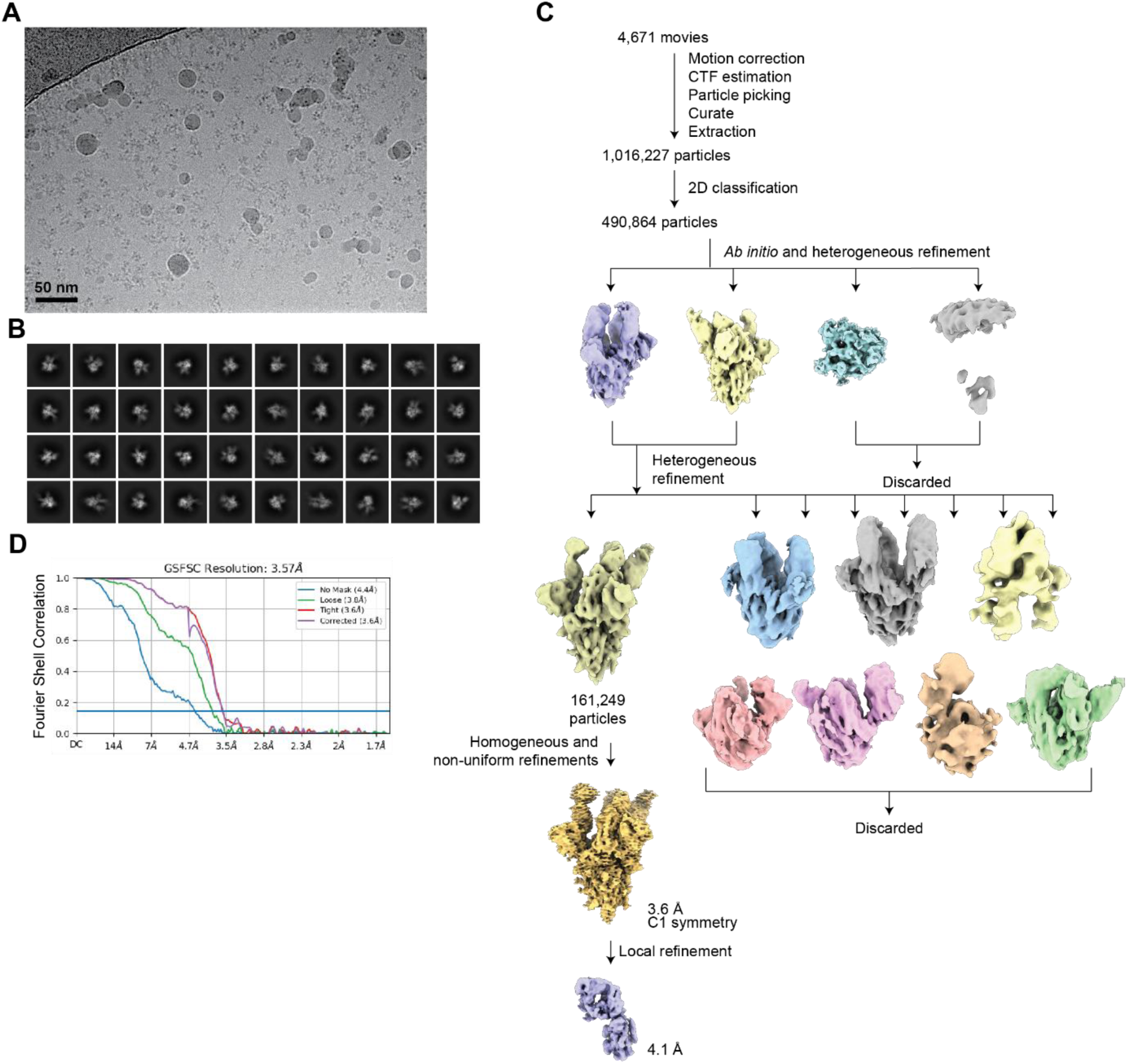
Cryo-EM data processing for 99 Fab-SARS-CoV-2 Lineage A spike complex. (A) Representative micrograph. (B) Representative 2D classes. (C) Workflow of single-particle cryo-EM data processing. (D) Fourier shell correlation (FSC) plots of the final reconstructions.

**Supplementary Figure 9.**
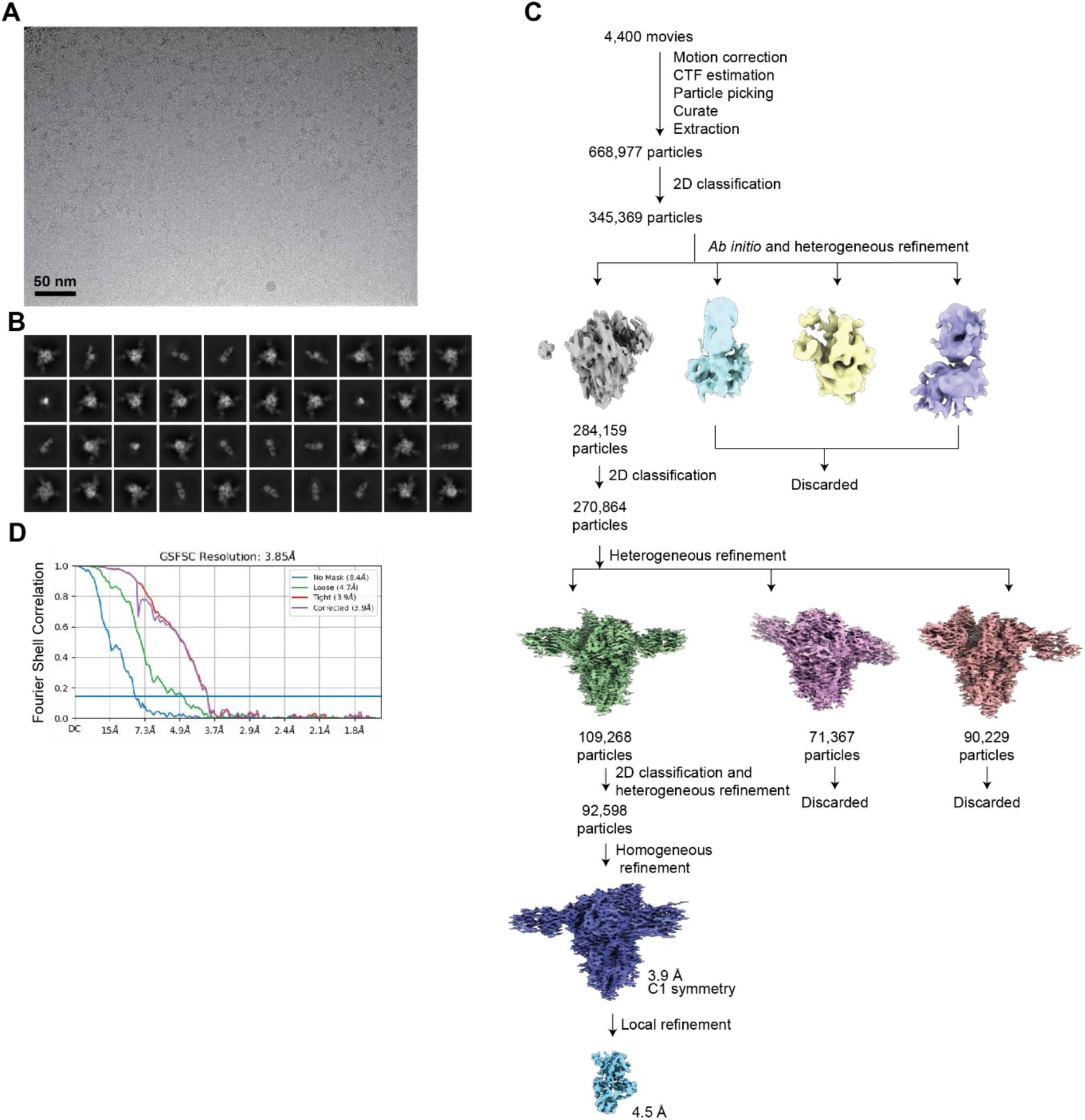
Cryo-EM data processing for 115 Fab-SARS-CoV-2 Lineage A spike complex. (A) Representative micrograph. (B) Representative 2D classes. (C) Workflow of single-particle cryo-EM data processing. (D) Fourier shell correlation (FSC) plots of the final reconstructions.

## Supplementary Tables

**Supplementary Table 1.**
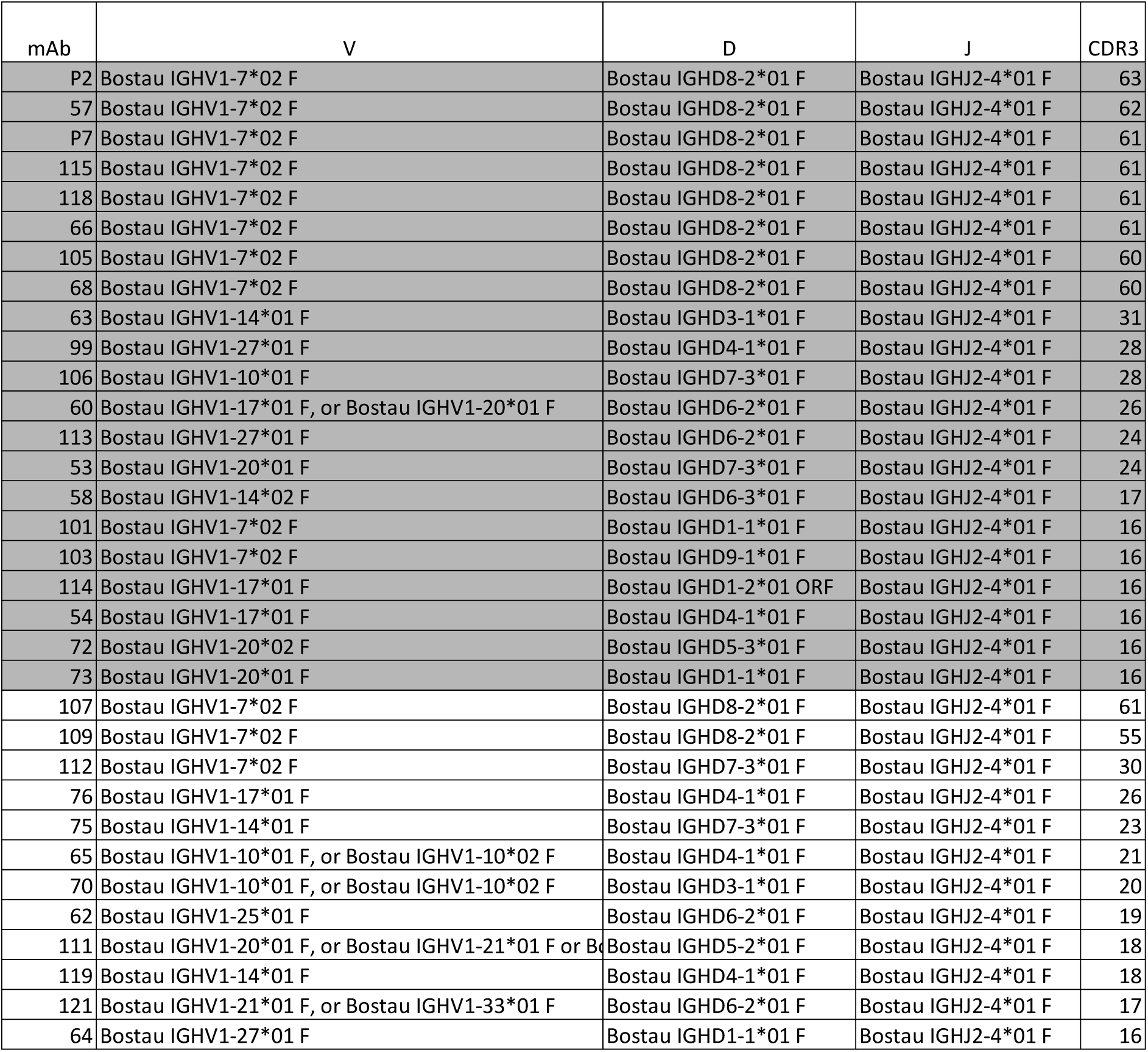
Table showing the VDJ genes and CDRH3 lengths of the mAb panel generated in this study. It consisted of 33 antibodies, and they are sorted based on the length of their CDRH3 length in amino acids. Grey (n=21) represents the spike binding mAbs whereas white shows the non-binders. Antibodies with CDRH3 >50aa were considered ultra-longs. VDJ germlines assigned by IMGT (69).

**Supplementary Table 2.** Histological changes and viral antigen expression in the lungs of male Syrian hamsters, aged 8-10 weeks that had received PBS (cohort 1), mAb 99 (cohort 2), mAb P7 (cohort 3) or Sotrovimab (cohort4) 24 h prior to intranasal infection with 10^4 PFU SARS-CoV-2 Delta in 200 μL and euthanised at 7 dpi.

| Animal ID | Treatment | Histological and immunohistological findings (lung) |
| --- | --- | --- |
| 1.1 | Cohort 1<br>[PBS IP, 24 h PI] | **HE:** multifocal consolidated areas with activated and hyperplastic type II pn, several NL and numerous mononuclear cells, and areas of BEC hyperplasia<br>**IH:** one pos macrophage in a consolidated area |
| 1.2 |  | **HE:** multifocal consolidated areas with activated and hyperplastic type II pn, some NL and numerous mononuclear cells, and areas of BEC hyperplasia<br>**IH:** one pos macrophage in a consolidated area and two very small patches of unaltered alveoli with a few pos AEC |
| 1.3 |  | **IH:** multifocal consolidated areas with activated and hyperplastic type II pn, several NL and numerous mononuclear cells, and areas of BEC hyperplasia; mild LC-dominated mononuclear pv infiltration<br>**IH:** one pos macrophage in a consolidated area |
| 1.4 |  | **Lungs:** multifocal consolidated areas with activated and hyperplastic type II pn, several NL and numerous mononuclear cells, and areas of BEC hyperplasia; mild LC-dominated mononuclear pv infiltration<br>**IH:** neg |
| 1.5 |  | **Lungs:** extensive multifocal consolidated areas with activated and hyperplastic type II pn, several NL and numerous mononuclear cells, and areas of BEC hyperplasia; mild LC-dominated mononuclear pv infiltration<br>**IH:** two consolidated areas with each one pos macrophage; alveolus with 2-3 pos AEC |
| 1.6 |  | **HE:** multifocal consolidated areas with activated and hyperplastic type II pn, several NL and numerous mononuclear cells, and areas of BEC hyperplasia; focal pb LC accumulation<br>**IH:** a few very small patches of pos macrophages and/or AEC, close to consolidated areas |
| 2.1 | Cohort 2<br>[mAb 99 IP, 24 h PI] | **Lungs:** NHAIR<br>**IH:** neg |
| 2.2 |  | **Lungs:** NHAIR, apart from minimal to mild mononuclear pv infiltration around a few small vessels<br>**IH:** neg |
| 2.3 |  | **Lungs:** NHAIR<br>**IH:** neg |
| 2.4 |  | **Lungs:** one small focal consolidated area of BEC hyperplasia, with a few embedded leukocytes<br>**IH:** neg |
| 2.5 |  | **Lungs:** two loosely consolidated areas with activated and hyperplastic type II pn and BEC hyperplasia, a few NL and numerous mononuclear cells; mild LC-dominated mononuclear pv infiltration<br>**IH:** neg |
| 2.6 |  | **Lungs:** NHAIR<br>**IH:** neg |
| 3.1 | Cohort 3<br>[mAb P7 IP, 24 h PI] | **HE:** several focal areas of consolidation with activated and hyperplastic type II pn and/or BEC hyperplasia, a few NL and numerous mononuclear cells<br>**IH:** neg |
| 3.2 |  | **HE:** small focal consolidated area comprised of BEC hyperplasia, with a few embedded leukocytes; small area with macrophages and a few NL in alveolar spaces<br>**IH:** neg |
| 3.3 |  | **HE:** several random, mainly small focal consolidated areas of BEC hyperplasia, with a few embedded leukocytes<br>**IH:** neg |
| 3.4 |  | **HE:** a few, mainly small random focal consolidated areas of BEC hyperplasia, with a few embedded leukocytes<br>**IH:** neg |
| 3.5 |  | **HE:** a few, mainly small random focal consolidated areas of BEC hyperplasia, with a few embedded leukocytes<br>**IH:** neg |
| 3.6 |  | **HE:** a few, mainly small random focal consolidated areas of BEC hyperplasia, with a few embedded leukocytes<br>**IH:** neg |
| 4.1 | Cohort 4<br>[Sotrovimab<br>IM,<br>24 h PI] | **HE:** NHAIR, apart from small focal mixed cellular infiltrate adjacent to a small vessel<br>**IH:** neg |
| 4.2 |  | **HE:** NHAIR, apart from a mild mixed cellular infiltrate around one larger vessel<br>**IH:** neg |
| 4.3 |  | **HE** NHAIR<br>**IH:** neg |
| 4.4 |  | **HE:** NHAIR, apart from minimal mononuclear pv infiltration around a few small vessels<br>**IH:** neg |
| 4.5 |  | **HE:** NHAIR, apart from minimal mononuclear pv infiltration around one small vessel<br>**IH:** neg |
| 4.6 |  | **HE:** NHAIR<br>**IH:** neg |

AEC – alveolar epithelial cells (type I and II pneumocytes); BEC – bronchiolar epithelial cells; HE – histological examination of hematoxylin-eosin (HE) stained sections from the left lungs; IH – immunohistology (IH) for SARS-CoV-2 nucleoprotein; IM – intramuscular; IP – intraperitoneal; LC – lymphocytes; neg – negative (no evidence of viral antigen expression); NHAIR – No histological abnormality is recognised. NL – neutrophils; PI – pre infection; pn – pneumocytes; pv – perivascular

**Supplementary Table 3.** Single-particle cryo-EM data collection, processing, and refinement.

| Structure | SARS-CoV-2 Lineage A Spike 6P + 99 Fab |
| --- | --- |
| **Data collection and processing** |  |
| Microscope | Caltech Titan Krios |
| Camera | Gatan K3 |
| Magnification | 105,000 |
| Voltage (keV) | 300 |
| Exposure (e<sup>-</sup>/Å<sup>2</sup>) | 60 |
| Pixel size (Å) | 0.832 |
| Defocus Range (μm) | -1 to -3 |
| Initial Particle Image (no.) | 490,864 |
| Final Particle Image (no.) | 161,249 |
| Symmetry Imposed | C1 |
| Map Resolution (Å) | 3.6 |
| FSC Threshold | 0.143 |
| **Refinement** |  |
| Initial Model Used | PDB 7UZ6, 6E9H and 8ECQ |
| Model Resolution (Å) | 2.8 |
| FSC Threshold | 0.143 |
| Model composition |  |
| non-hydrogen atoms | 26,057 |
| protein residues | 3,274 |
| ligands | 42 |
| Average B-factors (Å<sup>2</sup>) |  |
| protein | 107.9 |
| ligands | 109.3 |
| R.m.s. deviations |  |
| Bond length (Å) | 0.002 |
| Bond angles (°) | 0.571 |
| Validation |  |
| MolProbity score | 2.4 |
| Clashscore | 13.6 |
| Rotamer outliers | 3.7 |
| Ramachandran plot |  |
| Ramachandran favored (%) | 95.1 |
| Ramachandran allowed (%) | 4.9 |
| Ramachandran outliers (%) | 0 |
| PDB ID | 9DSL |
| EMDB ID | EMD-47146 |

**Supplementary Table 4.**
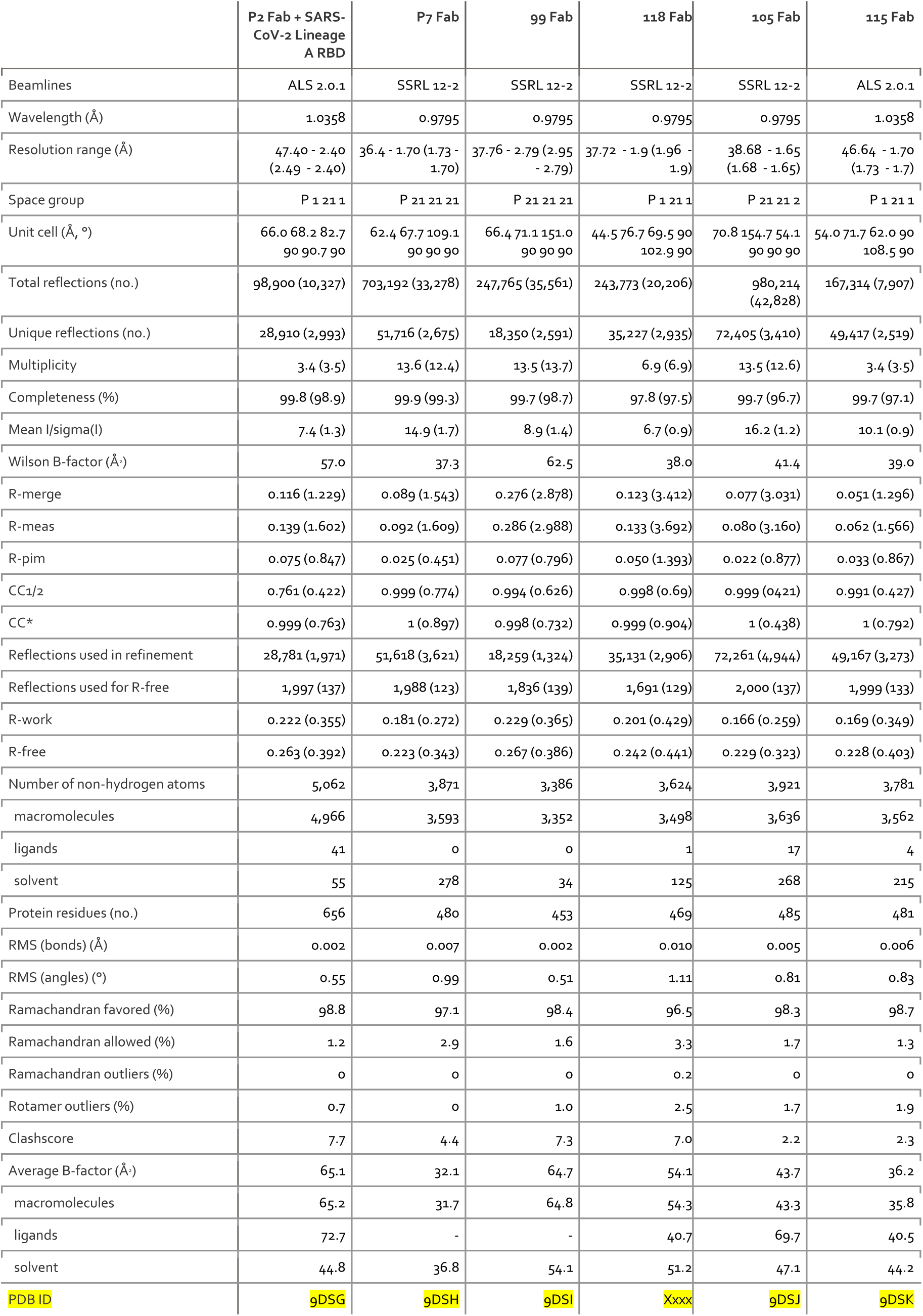
Crystal structure data collection and refinement.

## Notes

### Competing Interest Statement

The authors have declared no competing interest.

### Summary of Updates

Performed additional pseudovirus neutralization experiments with contemporary variants. Updated Figure 2B and added 2 authors.

